# Discovering genomic regions associated with the phenotypic differentiation of European local pig breeds

**DOI:** 10.1101/2022.02.22.481248

**Authors:** Klavdija Poklukar, Camille Mestre, Martin Škrlep, Marjeta Čandek-Potokar, Cristina Ovilo, Luca Fontanesi, Juliette Riquet, Samuele Bovo, Giuseppina Schiavo, Anisa Ribani, Maria Muñoz, Maurizio Gallo, Ricardo Bozzi, Rui Charneca, Raquel Quintanilla, Goran Kušec, Marie-José Mercat, Christoph Zimmer, Violeta Razmaite, Jose P. Araujo, Čedomir Radović, Radomir Savić, Danijel Karolyi, Bertrand Servin

## Abstract

**Background:** Intensive selection of modern pig breeds resulted in genetic improvement of productive traits while local pig breeds remained less performant. As they have been bred in extensive systems, they have adapted to specifical environmental conditions resulting in a rich genotypic and phenotypic diversity. This study is based on European local pig breeds genetically characterized using DNA-pool sequencing data and phenotypically characterized using breed level phenotypes related to stature, fatness, growth and reproductive performance traits. These data were analyzed using a dedicated approach to detect selection signatures linked to phenotypic traits in order to uncover potential candidate genes that may be under adaptation to specific environments.

**Results:** Genetic data analysis of European pig breeds revealed four main axes of genetic variation represented by Iberian and modern breeds (i.e. Large White, Landrace, and Duroc). In addition, breeds clustered according to their geographical origin, for example French Gascon and Basque breeds, Italian Apulo Calabrese and Casertana breeds, Spanish Iberian and Portuguese Alentejano breeds. Principal component analysis of phenotypic data distinguished between larger and leaner breeds with better growth potential and reproductive performance on one hand and breeds that were smaller, fatter, and had low growth and reproductive efficiency on the other hand. Linking selection signatures with phenotype identified 16 significant genomic regions associated with stature, 24 with fatness, 2 with growth and 192 with reproduction. Among them, several regions contained candidate genes with possible biological effect on stature, fatness, growth and reproduction performance traits. For example, strong associations were found for stature in two regions containing the *ANXA4* and *ANTXR1* genes, for fatness containing the *DNMT3A* and *POMC* genes and for reproductive performance containing the *HSD17B7* gene.

**Conclusions:** The present study on European local pig breeds used a dedicated approach for searching selection signatures supported by phenotypic data at the breed level to identify potential candidate genes that may have adapted to different living environments and production systems. Results can be useful to define conservation programs of local pig breeds.

## Background

In the last decades, pig breeding has focused mainly on improving growth rate, carcass leanness, and reproductive performances (1) of relatively few breeds (2). In parallel, most local breeds have not been subjected to such intensive management or genetic improvement and their use has declined. Local breeds are often raised in extensive farming systems, resulting in adaptation to specific environmental conditions and (often) poor feeding resources (3). However, this adaptation to seasonal fluctuations in feed availability may have resulted in low productivity (4). As a result, many local pig breeds have been abandoned or have even become extinct, while most of them have faced population bottlenecks and genetic drifts or introgression from other pig populations (5,6).

Today, many local breeds are used on a relatively limited scale and the available information on their phenotypic and genotypic traits is well known for only a few of them, such as the Iberian or the Meishan (7). Nonetheless, interest in local pig breeds has increased recently for several reasons, including their meat quality (allowing the production of high-quality meat products), their adaptation to local feeding resources, and society’s awareness of phenotypic and genetic biodiversity conservation (3). Being exposed to specific selection pressures in different local environments, local pig breeds also represent interesting genetic resources (8). They could become more important in the future as a reservoir of diversity useful to adapt to global change.

Recently, a genetic characterization of 20 European local pig breeds was carried out. It showed that some local breeds are clustered according to their geographical distribution (e.g. French Gascon and Basque breeds, Italian Apulo Calabrese and Casertana breeds, Spanish Iberian and Portuguese Alentejano breed), while some others suffer from introgressions or admixture with modern pig breeds (e.g. Lietuvos Baltosios Senojo Tipo and Lietuvos Vietiné with Large White and Landrace pigs; Mora Romagnola with Duroc pig) (8–10). Consequently, these breeds have developed particular heterogeneous phenotypic traits that could reflect specific genetic potential and adaptation to different production systems. As the measurement of phenotypic traits in pigs is to some extent standardised, it is possible to compare different local breeds. As shown in the study of Čandek-Potokar and Nieto (3) reviewing productive traits, not only do European local pig breeds differ from modern breeds, but they also exhibit an extensive variation among themselves.

To better understand the genetic basis for variation in phenotypic traits in local pig breeds, several genome-wide association studies focused on detecting possible associations of loci with different phenotypic attributes, such as morphological, production or meat quality traits (11–14). However, these studies worked with small sample sizes, increasing the risk to give false negative results due to low statistical power (15). Another approach to search for the association between genetic polymorphisms and phenotype is to look for genomic regions that have responded to selection (i.e., selection signatures). Several studies in local breeds have shown that the signals detected contain gene variants/genes that may be associated with variations in phenotype, such as coat colour, growth, reproduction, or fatness (8,9,16–19). The study by Muñoz et al. (9) determined the signatures of selection using SNP-array data in 20 European local breeds. This study revealed putative selection signals for regions containing genes involved in fatness, growth, reproduction, development, behaviour and sensory perception. To increase precision on the localization of selection signals, whole-genome sequencing was performed on DNA-pool samples from the same breeds/animals, and new analyses detected several regions associated with coat colour, body size, growth, reproduction and fat deposition (8). These two studies exploited methods based on the differences in allele frequencies between breeds to detect selection signals. Other approaches have been proposed, that jointly exploit information on allele frequencies and population-level phenotypes (20,21). They can offer better power to detect adaptations specific to some traits or environmental covariates. In the present study we adapt such an approach to combine the pool sequencing data from 19 European local and 3 modern pig breeds from (8) and the database of many phenotypic traits in 20 local pig breeds associated with stature, fatness, growth and reproductive performance from (3), in order to identify further selection signatures linked to specific breed level phenotypes and provide hypotheses on the physiological processes involved in the genetic divergence between local breeds.

## Methods

The aim of this study was to locate genomic regions associated with signatures of selection of local pig breeds for production traits. This was performed by combining genetic and phenotypic data at the breed (population) level. Most of the data used for this study have been previously published (3,8,9). In this section, genetic and phenotypic datasets are presented, and then the methodological approach to detect signatures of selection on phenotypic traits is described.

### Genetic Datasets

The present study was based on genetic data collected from 19 populations of European local pig breeds and 7 populations of industrial breeds. The data consisted of SNP genotyping performed with a medium density array (9) for 20 local breeds and whole genome sequencing of the pooled samples (8) for 19 of them (the Iberian breed was not included). To better describe the genetic structure of the local pig breeds, samples from additional 7 populations of 4 modern pig breeds were added. The final sample used for this study is shown in Table 1. Summary of whole-genome sequencing statistics was previously described in (8) and is presented also in Additional file 1, Table S1.

**Table 1.** Name of pig breeds, country of origin of the breeds.

| Pig breed name | Country | Number of<br>Individuals<br>genotyped on SNP<br>array | Sample size in<br>sequenced pool |
| --- | --- | --- | --- |
| **Local breeds** |  |  |  |
| Alentejano | Portugal | 48 | 35 |
| Apulo Calabrese | Italy | 53 | 35 |
| Basque | France | 39 | 30 |
| Bísaro | Portugal | 48 | 35 |
| Black Slavonian | Croatia | 52 | 35 |
| Cinta Senese | Italy | 51 | 35 |
| Gascon | France | 48 | 30 |
| Iberian | Spain | 48 | - |
| Krškopolje | Slovenia | 52 | 35 |
| Lietuvos Baltosios Senojo Tipo | Lithuania | 48 | 35 |
| Lietuvos Vietinė | Lithuania | 48 | 35 |
| Mangalitsa | Serbia | 50 | 35 |
| Mora Romagnola | Italy | 48 | 35 |
| Moravka | Serbia | 49 | 35 |
| Negre Mallorquí | Spain | 48 | 35 |
| Nero Casertano | Italy | 53 | 35 |
| Nero Siciliano | Italy | 48 | 35 |
| Sarda | Italy | 48 | 35 |
| Schwäbisch-Hällisches | Germany | 49 | 35 |
| Turopolje | Croatia | 49 | 35 |
| **Modern breeds** |  |  |  |
| Italian Large White | Italy | 4 | 35 |
| Large White | France | 97 | - |
| Italian Duroc | Italy | 5 | 35 |
| Duroc | France | 33 | - |
| Italian Landrace | Italy | 4 | 35 |
| Landrace | France | 53 | - |
| Pietrain | France | 61 | - |

### Quality control of genetic datasets

For the SNP array data, quality control of SNP genotypes was performed for the entire dataset using standard filters: only autosomal SNPs with < 10% missing data were retained. Following this step, 10 individuals with > 3% missing genotypes were clear outliers in the sample and were therefore discarded.

De novo SNV discovery was carried out on pool-sequencing data using CRISP (22) with default parameters and yielded 34,751,691 variants. From these, we filtered out variants with more than two alleles, variants with very low allele frequency that most likely resulted from sequencing errors (field VP = 0 and AF=1), and variants with low mapping quality (QUAL < 1000, MQ < 20). After filtering, 16,403,270 SNPs were kept. SNP allele frequencies were estimated for all discovered variants.

### Genetic structure of pig populations

The genetic structure of pig populations was assessed from individual SNP genotyping data using principal component analysis (PCA), admixture analysis (23) and population tree reconstruction with hapFLK (24) to reveal the genetic structure and covariance between populations. Admixture analysis was performed for a number of clusters (K) between 2 to 40 to determine a value of K that best explained the data. The decrease in cross-validation error was monotonous from 2 to 40 (Additional File 2: Figure S1), but showed a diminishing decrease after K=24, therefore this value was used as a reference to describe the genetic structure of the breeds. Based on the admixture results, individuals showing more than 80% of their genome assigned to the main cluster of their assigned breed were selected for inclusion in the population tree analysis. The Nero Siciliano breed did not show any individual matching to this criterion and was, therefore, not included in the population tree reconstruction.

To verify the quality of the pool sequence data, a comparison with the SNP array genotyping results was conducted. Allele frequencies of SNP array variants in the pooled sequence data were estimated using allele counts extracted with Samtools mpileup (25) and PoPoolation2 (26). Samtools was run with options −C 50 −q 20 (variants with mapping quality less than 20 were discarded as recommended by the software documentation). PoPoolation2 was run with default parameters on the resulting mpileup file. From the resulting pool allele counts, allele frequencies and Fst for each pair of populations were then calculated using the approach of (27) implemented in the R package poolfstat. The population tree was constructed applying the neighbour-joining algorithm on the Fst matrix using the same procedure that was used on allele frequencies derived from the SNP array genotypes.

### Phenotypic characterization of local pig populations

A database of phenotypic traits of European local pig populations (3) was used to determine global breed differences in phenotype. It gathers the results of different studies where the main experimental unit for the phenotypic trait was a trial, experiment or part of the experiment in which rearing conditions were sometimes very different from usual production conditions. However, all traits/variables considered here are standard measurements that can be directly compared across breeds and studies. Phenotypic variables were combined into four distinct groups summarizing stature, fatness, growth, and reproductive performance. The growth performance group included traits of average daily gain in 3 different growth periods, where the animals are fed *ad libitum* (i.e., from lactation to early fattening phase; up to approximately 60 kg). The stature group included traits of body weight and height of adult male and female animals. The fatness group comprised backfat thicknesses at different anatomical locations, fatty acid composition, carcass lean meat content and intramuscular fat content. The reproductive performance group included traits related to number of piglets per litter, number of litters per year, piglets/litter weights, and duration of lactation and farrowing interval. A more detailed description of the traits included in the phenotypic characterization can be found in Additional File 1: Table S2.

Means for each variable and each breed were calculated and scaled. Since the phenotypic database was composed of results from different studies, some variables were missing for some breeds (Additional File 2: Figure S4). Missing data were imputed using regularized iterative PCA method using R package missMDA (28). All experimental units extracted from all studies were given equal weight regardless of the number of pigs involved. Principal component analyses were performed using the R package FactoMiner (29) for the growth performance, reproductive performance, stature and fatness traits. Uncertainty in the predictions of missing data was assessed by multiple imputation (MIPCA function (30)). Breed loadings on the PC1 of each PCA were used as “breed scores” for the analysis of selection signatures. All statistical analyses were performed using R statistical software.

### Genome scan for selection on phenotypic breed scores

To identify genomic regions associated to selection on breed level phenotypes, we built upon the approach of (20) which we briefly describe here. Assuming data on allele frequencies measured in r populations at L loci, this association model at a locus l is:

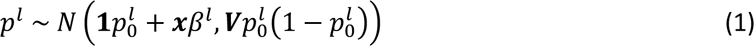

where ***p****^l^* is the vector of allele frequencies in the r populations, 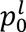 is the (unknown) ancestral allele frequency of the locus, ***x*** is the vector of breed level phenotypes, **β***^l^* is the effect of phenotype on allele frequencies differences, and ***V*** is the genome-wide variance-covariance matrix of allele frequencies between populations. The idea behind this model is that the adaptation of populations to covariate *x* (here the breed level phenotypes) will drive allele frequency differences between populations away from their expectation under genetic drift. In (1), this expectation is modelled with the genome-wide covariance matrix **V** (see (20) for the detailed description). To perform statistical inference under this model the parameters **V**, β*^l^* and 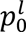 need to be estimated. Under the null hypothesis (selection associated with phenotype x has not affected allele frequencies), the parameter β*^l^* is set to 0. To test for association with the covariate **x**, (20) use a MCMC algorithm allowing to derive a statistic for association (a Bayes Factor). This was later extended by (21) to account for uncertainty in allele frequency estimation in pool sequencing experiments. This approach was tested on our dataset, but was found computationally inefficient due to the very high number of SNPs considered. We therefore used a frequentist treatment of the model that consists in maximizing the likelihood of model (1) under the null and the alternative hypothesis and performing a likelihood ratio test. One deviation from this approach is that we account for the variation in sequencing depth in different pools by using regularized allele frequencies 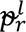 (see below) rather than fitting the model on the usual allele frequency estimates 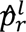. The maximum likelihood estimator of allele frequencies is 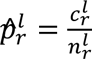, where 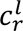 is the number of alternative allele counts and 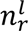 the sequencing depth for population r at locus l. This estimator has good properties provided the sequencing depth 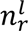 at the locus is large. However, this depth is highly variable along the genome and can be quite low (even 0) in some genomic regions. Moreover, it will vary between populations at a given locus. To regularize allele frequencies estimates, we used shrunk allele frequencies estimates in the form:

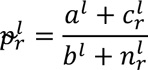

where *a^l^*and *b*^l^are regularizing (prior) parameters set so that *a^l^*/*b^l^* equals the alternative allele frequency among all populations at locus l. Doing so, if a population has no observed data at a SNP, its allele frequency will be similar to other populations which reduces the risk of false positives due to uneven sequencing coverage.

The association of breed scores to allele frequencies was tested for all SNPs for all breed scores (stature, growth performance, fatness traits, reproductive performance). The result of the test was an asymptotic p-value for each SNP for each phenotypic trait. Since model (1) is only valid for intermediate ancestral allele frequencies, only SNPs where the ancestral allele frequency estimate 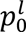 lied between 0.05 and 0.95 were considered in the following. To reduce the possibility of false positives, a permutation procedure was conducted to correct the asymptotic p-values. For each phenotype tested, a new analysis was performed on a permutation of the phenotype, *i.e.* under the null hypothesis of no association between the phenotype and the allele frequencies. This produced an empirical distribution of the asymptotic p-values under the null that was used to obtain corrected p-values at each SNP. Statistical significance was then established on the corrected p-values by estimating the False Discovery Rates (FDR) with the approach of (31) using the qvalue R package. The FDR threshold for detecting significant associations was set at 1%.

Based on association statistics for individual SNP, we defined association regions and extracted candidate genes from the annotation of the reference genome (*Sscrofa11.1*). Regions containing at least 4 significant SNPs less than 200Kb apart were further annotated. The detected genes within these regions were reviewed in the literature for potential biological effects on phenotypic differentiation.

## Results

### Genetic structure of European pig breeds

To characterize the genetic structure of populations, the individual genotypes on the SNP array were used to perform a standard genetic structure analysis. PCA analyses of 24 breeds revealed main genetic backgrounds in the dataset (i.e. Iberian, Duroc/Mora Romagnola, Large White and Landrace/Pietrain backgrounds), visible in the first 3 principal components (Figure 1). PC1 and PC2 clearly distinguish between the Iberian pig, White pigs (Landrace, Large White and Pietrain) and Duroc/Mora Romagnola backgrounds. PC3 shows that the White pigs’ background separates into a Large White specific background and a Landrace/Pietrain background. PC4 further separates the Turopolje breed from the Iberian group. The global pattern of differentiation between breeds in our dataset is thus strongly influenced by modern pig breeds. Local pig breeds are usually in intermediate positions, mostly within a triangle with summits corresponding to the Iberian breed, the Landrace/Pietrain and the Large White breeds (see Additional files 3). The two exceptions are the Mora Romagnola and Turopolje breeds, which appear much more differentiated. In order to interpret these patterns, further analyses using admixture clustering and population tree reconstruction were performed.

**Figure 1.**
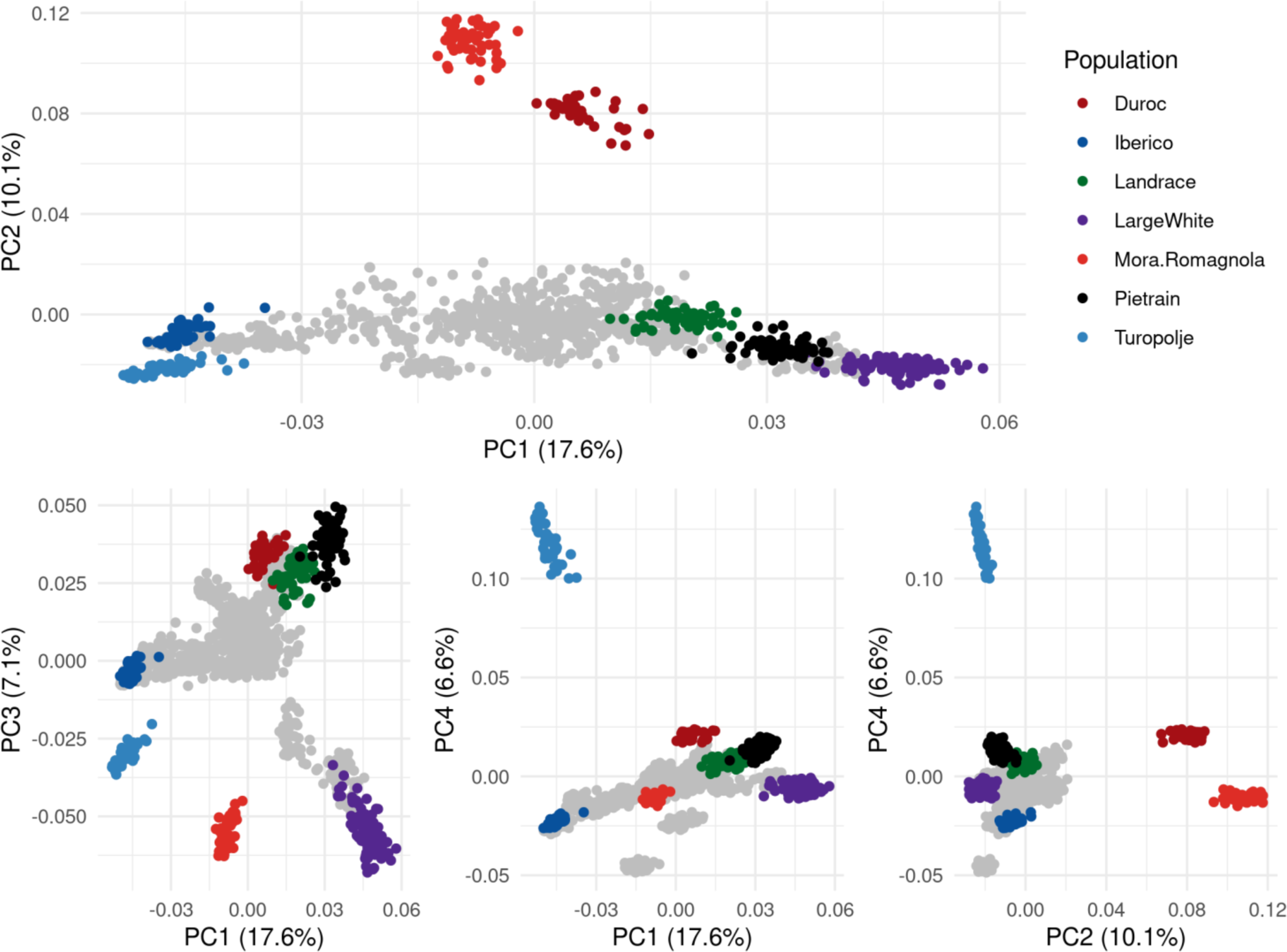
Principal component analysis of 24 pig breeds genotyped on a medium density SNP array.

The results of the admixture analysis results show that the number of homogeneous clusters in the dataset is difficult to determine. The cross-validation procedure was conducted for up to 40 clusters and resulted in a general decrease in cross-validation error with increasing number of clusters (Additional File 2: Figure S1). However, the decrease slowed down after K=24 clusters, a number that corresponds to the number of named breeds (Table 1). The further decrease is due to the fact that some breeds are sub-structured and require additional clusters to be well fitted. This is typical of local breeds (e.g Gama et al., 2013; (32)). In the following, we present the results obtained with K = 24 corresponding to the inflexion point in the cross-validation curve. Figure 2 shows side-by-side the result of the population tree reconstruction and the admixture analysis. At K=24, most breed exhibit a homogeneous pattern of admixture and belong to a specific cluster. Exceptions are the Alentejano and Iberian breeds (which belong to the same cluster, in consistence with their common origin), the Casertana and Apulo Calabrese breeds (each of them further splits into two groups) and Sarda, Moravka, Bisaro and Nero Siciliano breeds that show high heterogeneity. In the latter breeds, admixed individuals do not appear to be recent hybrids with other breeds. In addition, admixture plots with K = 20, K = 15, K = 10 and K = 6 are shown using circular plot in Additional file S2, Figure S2.

**Figure 2.**
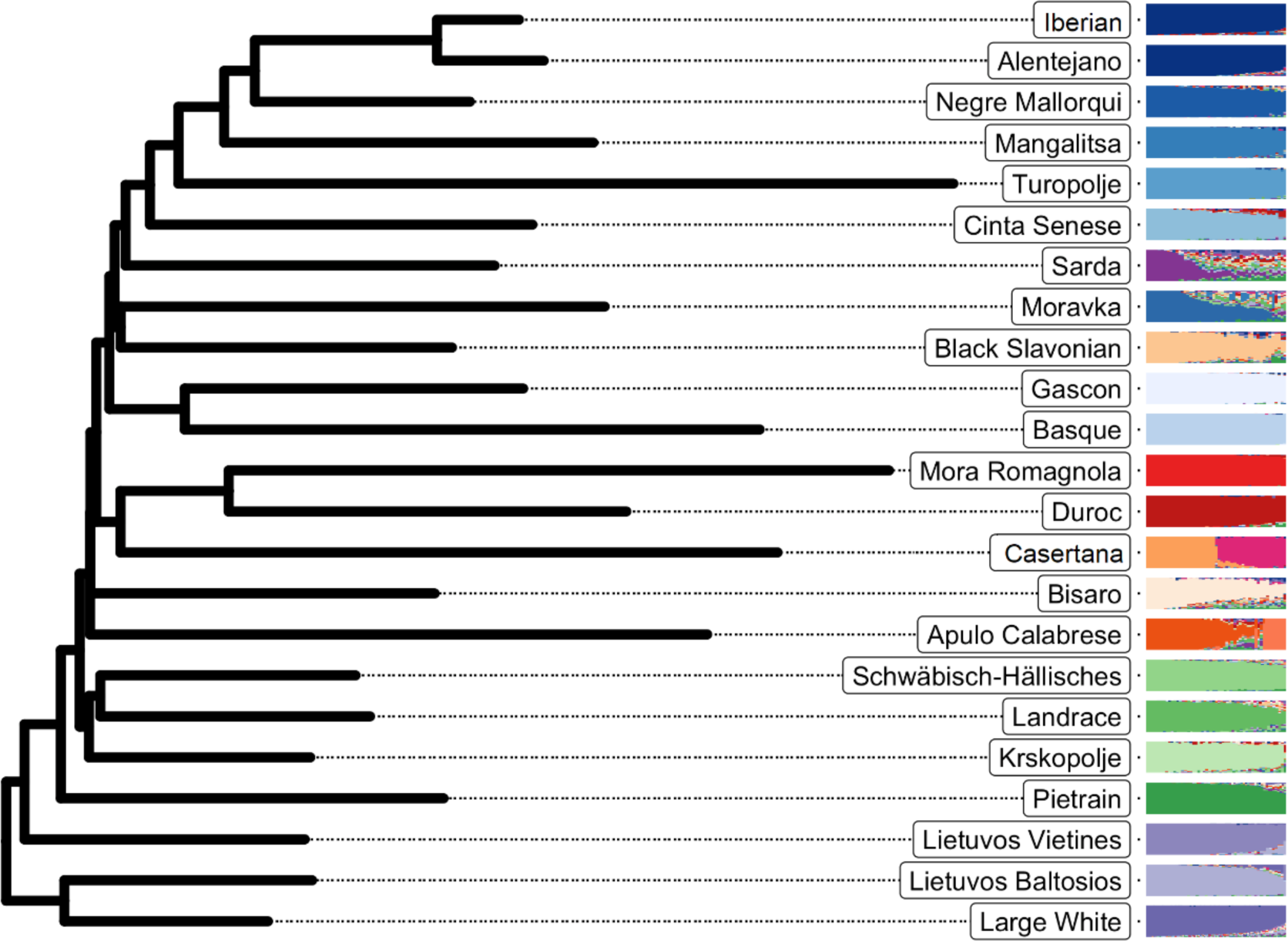
Population tree and admixture analysis of 23 pig breeds genotyped on a medium density SNP array. The population tree reconstructed from pairwise genetic distances (Fst) is shown on the left, and the admixture component for all individuals a priori belonging to the breed is shown on the right. The colour levels follow the global axes of genetic variation, with populations most closely related to the Iberian type shown in shades of blue, to the Duroc in shades of red, to the Large White in shades of purple, and to the Landrace in shades of green. Heterogeneous populations, or those equidistant from these four clusters, are shown in orange. The Nero Siciliano breed exhibited extremely high heterogeneity, and was therefore not included in the reconstruction of the population tree and is not shown in this figure.

The population tree is structured consistently with the main axes of variation identified in the PCA analysis with main genetic backgrounds (Figure 2). Turopolje and Mora Romagnola breeds differentiated early in the PCA analysis, which could be explained with the population analysis and the fact that they exhibit long branches corresponding to low heterozygosity. This analysis also reveals a general clustering of local breeds according to their geographical origin. Breeds closer to the Iberian group mostly originate from the Iberian peninsula or close geographical areas (South West of France with Basque and Gascon breeds, Balearic Islands with Negre Mallorqui breed). Interestingly, some other breeds from different geographical area appear to be related to this background such as Mangalitsa and Moravka from Serbia, the Black Slavonian from Croatia or the Cinta Senese from Italy. Breeds from Central Europe, such as Schwabisch-Hällisches and Krškopolje pig, showed genetic proximity to the Landrace/Pietrain background and breeds from Lithuania in Northern Europe with the Large White component. Finally, some breeds such as the Bísaro from Portugal or Apulo Calabrese from Italy cannot be considered related to any other breed in the dataset.

Sequences from the investigated breeds were aligned to the Sscrofa11.1 reference genome and SNPs were discovered with the pool-seq variant caller CRISP. For SNPs present on the genotyping array, allele frequencies were consistent with those derived from individual genotyping (9) (Additional file 2, Figure S3). This confirmed the quality of SNPs obtained by pool-sequencing.

### Phenotypic differentiation of European local pig breeds

A database of available published results on phenotypic traits of 20 local pig breeds (3) was used in order to distribute local pig populations according to phenotype including stature, growth, fatness traits, and reproductive performance traits. The representation of relationships between breeds and variables was evaluated using PCA analyses. The first two principal components of the PCA for stature, growth, fatness, and reproduction group accounted for 98.2%, 83.2%, 84.3% and 70.7% of the total variance, respectively (Figure 3, Additional File 1: Figures S5, S6, S7). Scores for each breed from PC of all phenotypic groups were extracted from PCA.

**Figure 3.**
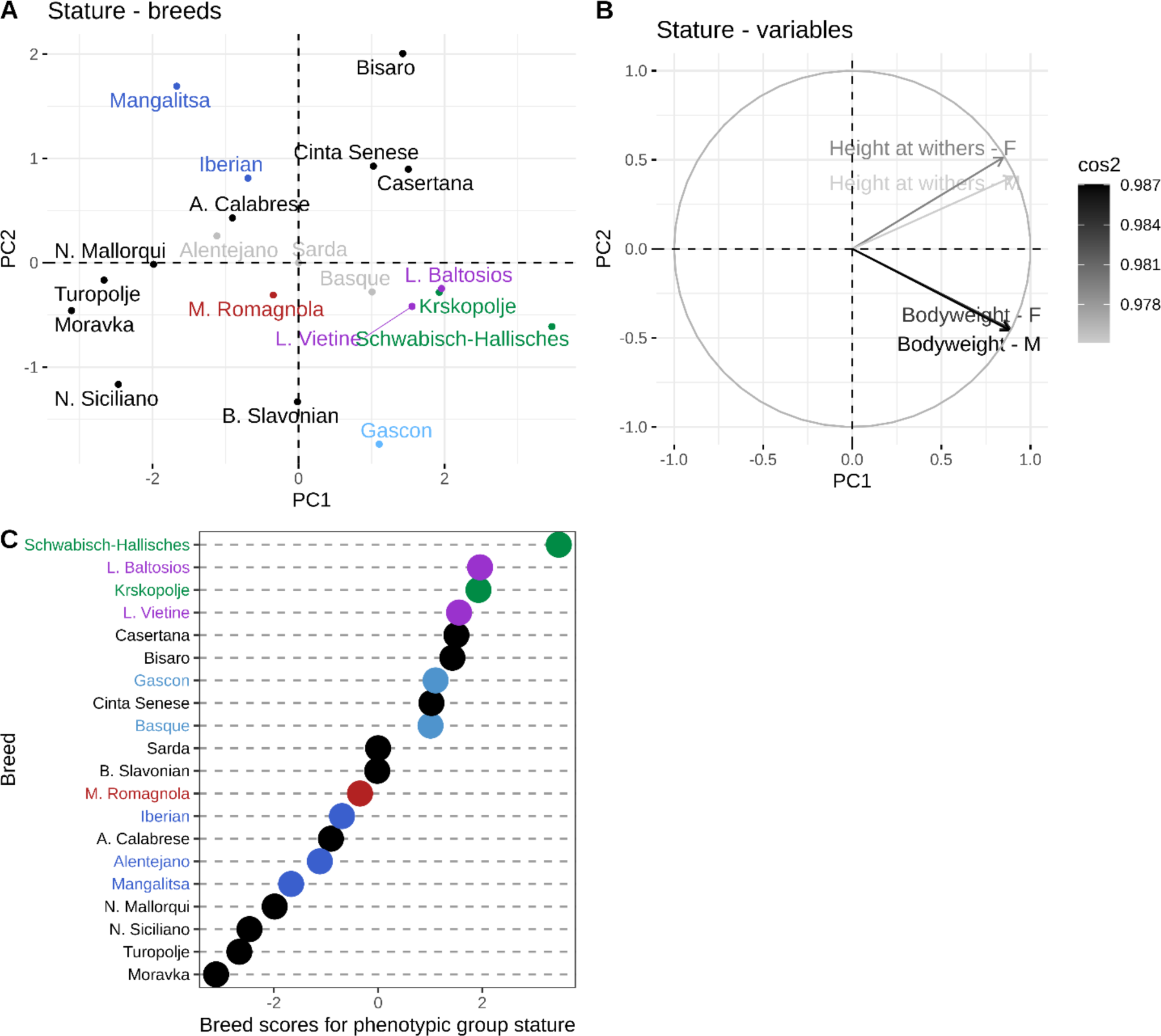
Principal component analysis showing the relationship between breeds (A) and the traits associated with stature (B) and the corresponding phenotypic breed scores (C). Breeds (A) coloured in grey are the breeds with more than 50% of missing variables, thus, their position on the PCA must be interpreted carefully. The variables (B) are coloured according to quality of the representation, which is measured by squared cosine between the vector originating from the element and its projection on the axis. The variables that contribute most to the separation of the trait into PC1 and PC2 are coloured black. Breeds (A, C) are coloured according to genetic similarity. Breeds (A, C) in green are genetically Landrace-like breeds, in purple are Large White-like breeds, in blue are Iberian-like breeds, in red are Duroc-like breeds and in light blue are Gascon and Basque.

The first principal component of the stature group (Figure 3) represented the majority of the total variability (77.3%) and clearly distributed breeds according to average height and weight. Thus, breeds with lower breed scores for stature (e.g. Moravka, Turopolje, Nero Siciliano, Negre Mallorqui) were smaller and lighter, while breeds with higher breed scores were taller and heavier breeds (e.g. Schwäbisch-Hällisches, Lietuvos Baltosios Senojo Tipo, Krškopolje pig, Lietuvos Vietiné).

The growth group distributed local breeds according to their growth capacity, including data on average daily gain for the rearing period from the lactation period to the early fattening phase (i.e. up to 60 kg body weight) (Additional File 2: Figure S5). The first PC explained 56.1% of the total variability and distributed the breeds with the highest (i.e. Schwäbisch-Hällisches, Lietuvos Baltosios Senojo Tipo, Lietuvos Vietiné, Bisaro, Krškopolje pig) and the lowest (i.e. Alentejano, Moravka, Black Slavonian, Mangalitsa and Turopolje) growth potential.

Fatness traits comprised variables associated with the fatty phenotype (Additional File 2: Figure S6). Principal component 1 (59.3% of total variability) was positively correlated with backfat thicknesses at different anatomical locations and intramuscular fat content. Conversely, lean meat and PUFA content (i.e., lean phenotype) were negatively correlated with PC1. Because intramuscular fat content is a trait of particular interest, the distribution of breeds by intramuscular fat content is shown in Additional File 2: Figure S8. Local breeds were divided into fatter (e.g. Moravka, Iberian, Mangalitsa, Negre Mallorqui) or leaner phenotypes (e.g. Schwäbisch-Hällisches, Lietuvos Baltosios Senojo Tipo, Bisaro).

The final characterization of local pig breeds was done according to reproductive performance (Additional File 2: Figure S7). Herein, the PC1 clearly distinguished breeds with larger litter sizes and piglet birth weights (i.e. Schwäbisch-Hällisches, Lietuvos Baltosios Senojo Tipo, Lietuvos Vietiné, Bisaro, Krškopolje pig) from breeds that had lower reproductive performance with smaller litters and lighter piglet birth weights (i.e. Nero Siciliano, Turopolje, Casertana, Mora Romagnola). Lastly, breed scores of phenotypic trait groups were plotted to examine the global distribution of breeds and production traits in local pig populations (Figure 4). Figure 4 shows that breeds characterised by larger size and higher growth potential were also more reproductively efficient than smaller breeds with lower growth rate. In addition, fatter breeds were smaller and lighter with smaller growth rate than leaner breeds that are larger.

**Figure 4.**
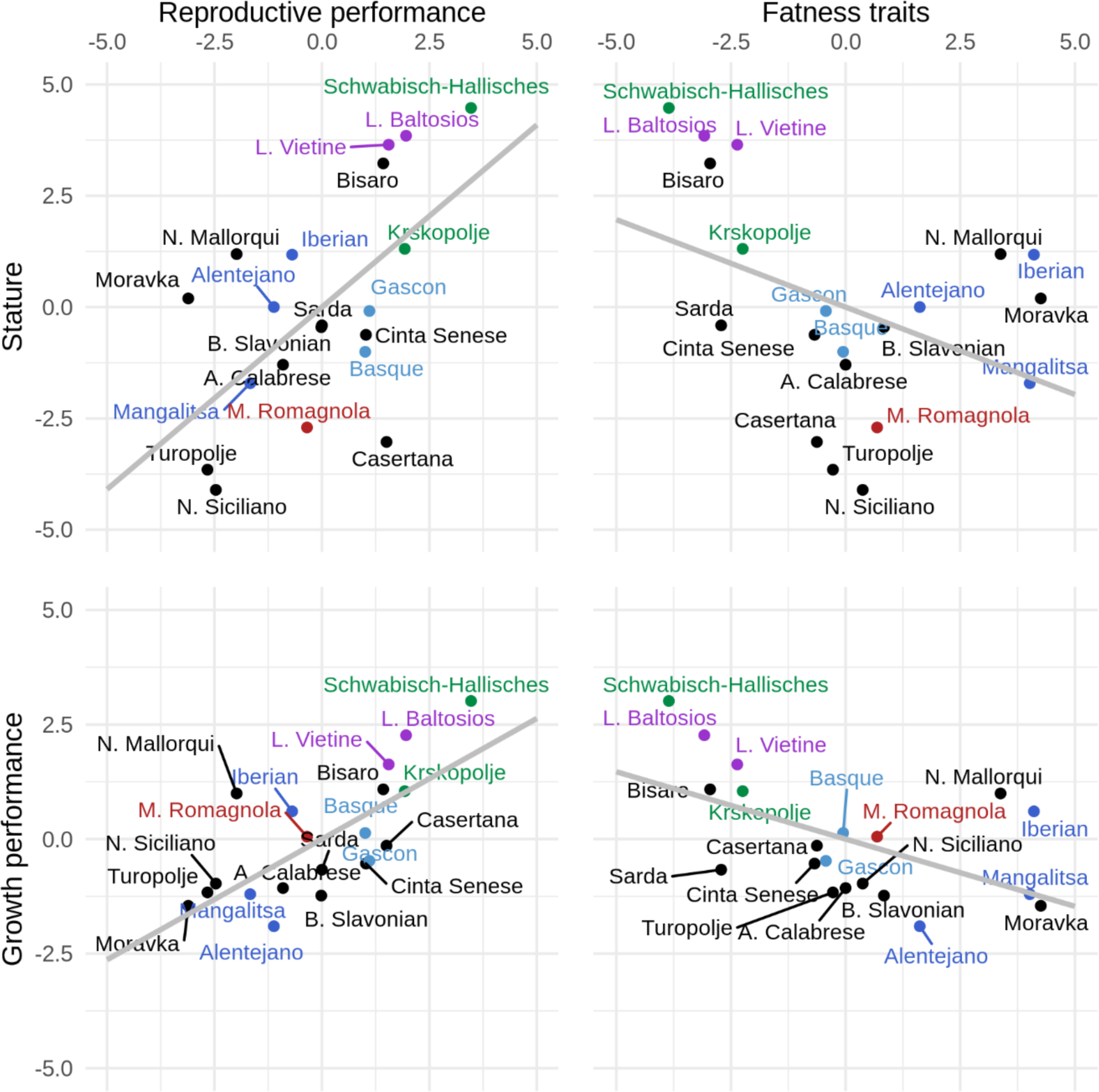
Global differences in production traits for 20 European local pig populations. Breeds are distributed according to breed scores for phenotypic traits and colored according to genetic similarity. Lower values for breed score growth are representing lower average daily gain from lactation up to 60 kg of live weight, while higher values represent higher average daily gain in the same growth period. Low values for breed score reproduction represent low reproductive performance, while high values represent higher reproductive performance of breeds. Low values for breed score stature represent lighter and smaller breeds, while high values represent heavier and larger breeds. Low values for the breed score fat represent leaner breeds, while high values represent fatter breeds. Breeds in green are genetically Landrace-like breeds, in purple are Large White-like breeds, in blue are Iberian-like breeds, in red are Duroc-like breeds and in light blue are Gascon and Basque.

### Detection of genomic regions associated with phenotypic differentiation

The approach proposed by Coop et al. (20) was extended to breed-level phenotypes to find genomic regions potentially influencing phenotypic differentiation in local European pig breeds (see Methods). A selection signature scan was performed for each phenotypic breed score, resulting in p-values for each SNP. Figure 5 shows Manhattan plots for stature, fatness, growth and reproduction along with a pp-plot constrasting the expected distribution of (-log10) p-values to the observed one.

**Figure 5.**
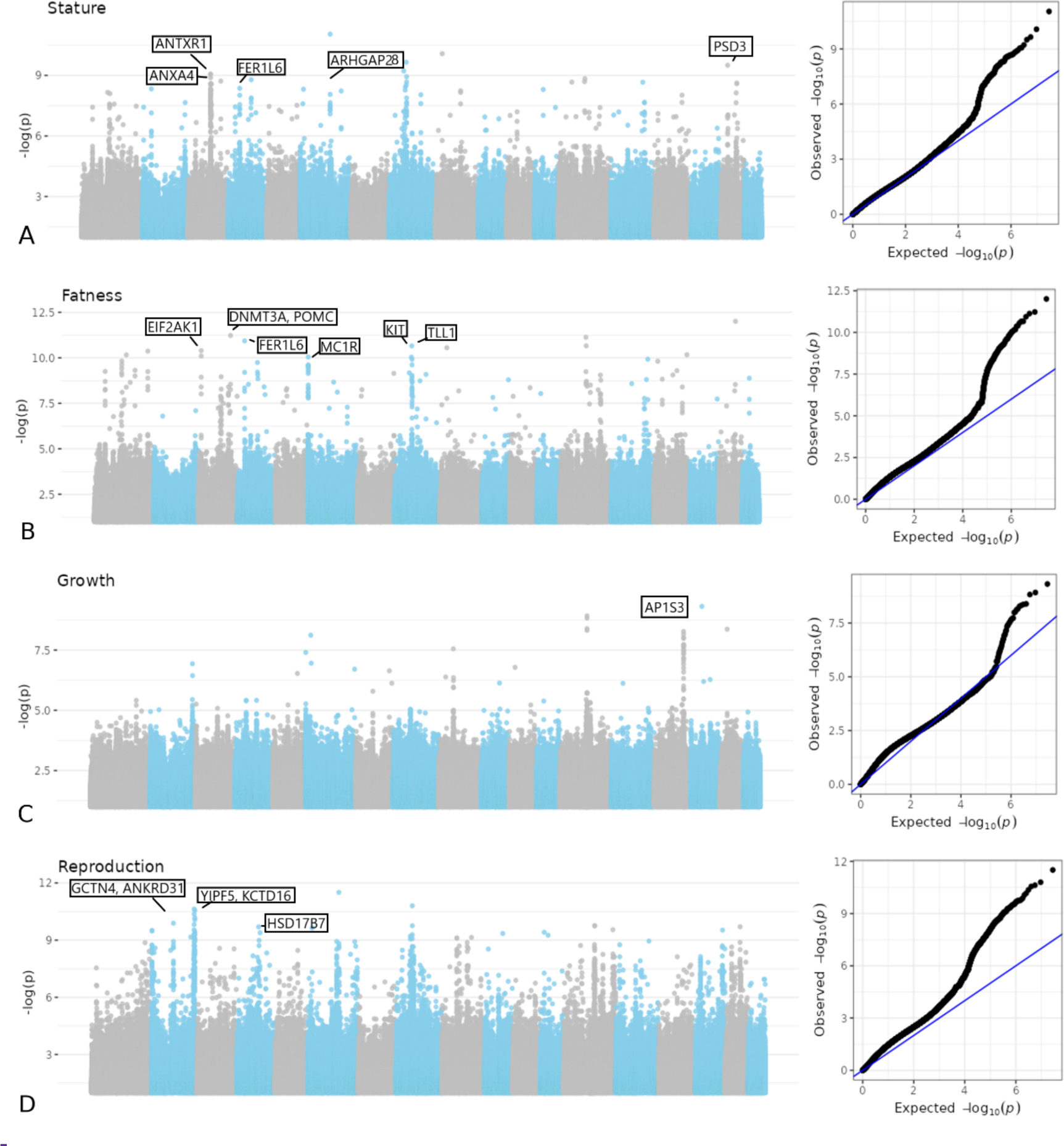
Manhattan plots for phenotypes (A) stature, (B) fatness, (C) growth and (D) reproduction and associated pp-plots.

Across the genome, windows/regions with 4 significant SNPs with significance threshold 0.01 were considered as selection signals for phenotypes. Overall, a total of 234 regions/windows ranging in length from 3.1 kbp to 1.282 Mbp were discovered. Among the detected regions, 16 regions were in the stature group, 2 regions were in the growth group, 24 regions were in the fatness group, and 192 regions were in the reproduction group. For a list of selection signatures for phenotypes, see Additional File 4.

Regions in the stature phenotypic group spanned between 3.1 kbp and 362.2 kbp with a maximum of 35 SNPs and included from none to four genes. The region with the strongest signal included the *ARHGAP28* gene. The growth phenotypic group contained 2 regions with a maximum of 18 significant SNPs and included 1 gene (i.e. *AP1S3*). The fatness phenotypic group spanned from 3.1 kbp to 590.5 kbp in length with maximum of 30 SNPs and included between up to 12 genes. The region with the strongest signal included the *DNMT3A, POMC* and *EFR3B* genes. The reproduction phenotypic group contained regions spanning from 12.8 kbp to 1,282.0 kbp in length with a maximum of 170 SNPs. The region with the strongest signals was located on chromosome SSC6 and contained the *YES1*, *ENOSF1*, *TYMS*, and *CLUL1* genes. The detected selection signals for production traits were further examined to find overlaps between groups (Additional file 2: Figure S9).

For each phenotypic group, we estimated the enrichment/depletion of significant SNPs in different functional categories: Intergenic, upstream of genes, genic (coding regions or introns) and downstream of genes (Figure 6). We observed a general trend for an enrichment of significant SNPs in genic regions (exons and introns) and some depletion in intergenic regions.

**Figure 6.**
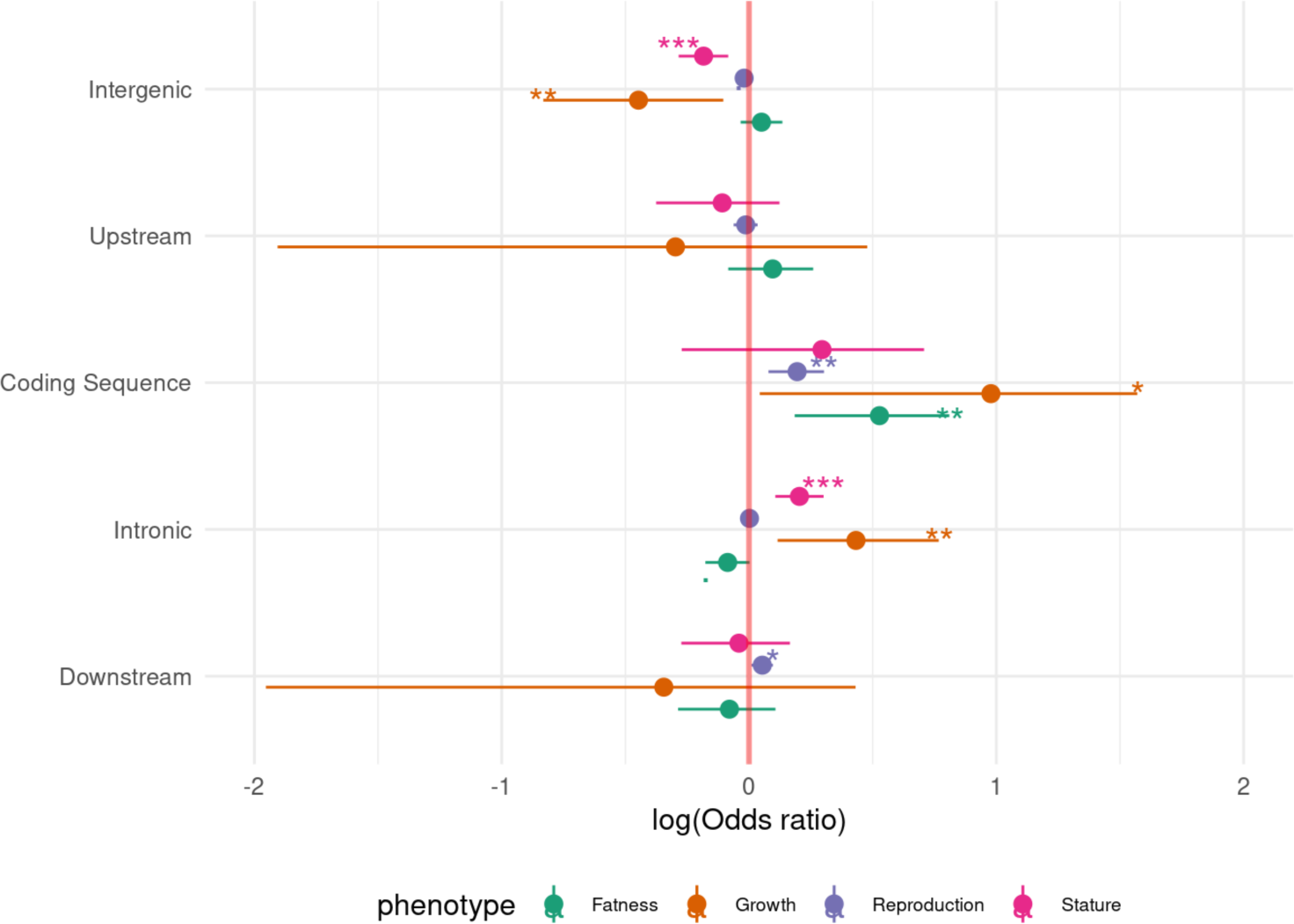
Enrichment of SNPs associated with different traits into functional categories.

## Discussion

Deciphering the genetic basis of variation in complex traits can help to understand their biology and evolution, and improve breeding, selection programs and conservation plans of animal genetic resources. One approach to address this question is to link genetic and phenotypic variation by identifying genomic regions that have evolved in response to selection on complex traits. Establishing such links requires a collection of data on genetic groups (e.g. populations, lines, breeds) that share a common origin and have information on adaptive traits that can be phenotyped in a standardized fashion. Livestock species generally meet these criteria and therefore provide good models for mapping genomic regions associated with selective constraints. Here, we studied local pig populations that were genetically characterized using individual genotyping and pool sequencing for which a recent common phenotypic database was constituted.

### Genetic diversity of European local pig breeds

Using individual genotyping of a large set of European breeds, we have found that their genetic variation is structured around four main genetic groups: the Duroc, the Large White, the Landrace and the Iberian. The origin of the Duroc is unclear, as it originated from American continent and was then brought back to Europe. This history implies that it was separated from the European breeds for a long time, which is evident in our genetic analysis by its rather large genetic distance with most European breeds, except for Mora Romagnola breed from Italy, which has a history of crossbreeding with Duroc breed (33). In addition, the present study showed that Mora Romagnola (and also Turopolje pig breed) differentiated early in the PCA analysis, which can be explained by their high degrees of inbreeding (34). The White pig group is composed of the Lietuvos breeds and the Large White, all of which originate from Northern Europe and are clearly separated from the Landrace, which includes populations originating from more central countries such as Germany (Landrace, Schwäbisch-Hällisches), Belgium (Pietrain) and Slovenia (Krškopolje). The Large White and Landrace groups are close to the root of the population tree, which could be explained by the well-documented influence of Asian pigs on these genetic pools (35). Finally, the Iberian group includes a set of breeds that are quite geographically dispersed; while many of the Iberian-type breeds are found around southwestern Europe (Spain, Portugal, and France), some populations from Eastern/Central Europe clearly belong to the same genetic pool (e.g. Mangalitsa from Serbia, Turopolje from Croatia, Cinta Senese from Italy). A possible interpretation is that the genetic background responsible for variation in the Iberian pig breed was historically widespread in Europe and can still be found in some local pig breeds. Overall, analysis of the genetic structure of European local pig breeds reveals extensive genetic variation clustered in differentiated breeds with continuously distributed genetic distances. The historical events that led to the present genetic variation are most likely complex, and involve differentiation from an ancestral pool followed by or accompanied by outcrossing between populations, including wild boars, and cannot be reconstructed from the data used here. However, the sample of available breeds is well adapted to our main objective, as we can interrogate a diverse set of polymorphisms for adaptation to contrasting environments or production systems in different genetic backgrounds.

While SNP genotyping is sufficient to characterize the genetic relationships and population structure of local pig breeds, the density of the SNP array used limits the resolution and power to detect potential associations with breed phenotypes especially in local breeds where a big proportion of SNPs included in the commercial chips are not segregating. To alleviate this issue, a cost-effective alternative to SNP genotyping consists of pool sequencing of populations. In the dataset used here, we confirmed that allele frequency and genetic structure estimates obtained from pool sequencing are consistent with those obtained from SNP array, while providing comprehensive information on the genetic diversity of local pig breeds.

### Phenotypic diversity of European local pig breeds

In parallel to the comprehensive information on the genetic diversity of local pig breeds, we developed an original approach based on PCA to characterize their phenotypic diversity in traits related to stature, growth, fatness and reproduction, which have been described in a comprehensive analysis (3). While local breeds can be phenotypically characterized on meat quality traits (pH24, pH48, intramuscular fat content and *longissimus dorsi* muscle colour, Additional File S2: Figure S10)., these traits (with the exception of IMF) are strongly influenced by pre-slaughter handling stress (36). Breed score for meat quality was therefore not included in the phenotype-genotype selection scan.

Local pig breeds exhibit a wide variety of exterior phenotypic characteristics, including body size or body weight. This could be due to genetic factors or could reflect the management of the breed. For example, male and female animals of the Schwäbisch-Hällisches breed are medium to large in size and also have well-developed management systems. In contrast, more untapped breeds (e.g. Moravka) are smaller and lighter (3).

Regarding the growth performance of local pig breeds, existing knowledge is limited. Moreover, the production systems may not be sufficiently adapted to the needs of the breeds which could be reflected in the phenotype of the breeds. For example, the study of Brossard et al. (37) argues that in many studies on local pig breeds, animals are fed below ad libitum level and, therefore, do not reach their full production potential. This is consistent with an analytical literature review (38) on European local pig breeds, which demonstrated that local pig breeds are generally fed in ad libitum conditions only earlier in life (during the growing and early fattening phase). In the late fattening phase, the feeding is likely to be restricted. To limit the influence of such effects in the present study and to use a better representation of breeds’ growth potential, only average daily gain values from the beginning of the lactation to the end of the early fattening phase (i.e. up to 60 kg) were used for the phenotypic score on growth. Despite this, better growth performance might still partly reflect breeding management, as shown for the Schwäbisch-Hällisches pig, which has a growth potential comparable to genetically improved modern pig breeds (39,40).

It has been reported that local pig breeds have a lower protein and higher lipid deposition capacity compared to modern pig breeds (37). The higher genetic capacity of local pig breeds for backfat and intramuscular fat deposition, and lower muscle accretion has been demonstrated in several studies comparing fat characteristics between local and modern breeds (41–45). Although the local pig breeds are recognised as fatty, they still exhibit a large phenotypic variability in the dataset used here.

Regarding reproduction traits, sows of local breeds typically exhibit relatively high age at first parturition, few litters per sow per year, long lactation periods, small litter sizes and high piglet mortality (38). In our study, PCA of reproductive performance distinguished breeds with better reproductive efficiency, which are usually reared in more intensive systems within more developed pork chains (e.g. Schwäbisch-Hällisches pig) from breeds with lower reproductive efficiency (e.g. Nero Siciliano, Turopolje, Casertana, Mora Romagnola, Mangalitsa) reared in extensive or semi-extensive systems (3).

While some of this variation in phenotypes can certainly be explained by differences in production systems among local breeds, our genetic association results suggest that some of it may be linked to genetic variation. In order to be able to test for these genetic associations, the methods used require all breeds to have an associated phenotype. To do so, we imputed using PCA the missing phenotypes of some breeds (see Methods). The resulting imputed phenotypes are likely to be biased toward the average of all breeds. This is for example most likely the case for the Sarda breed, considered to be a very small breed (in term of body size) but lying at an intermediate value after imputation. This certainly limits our power to detect genetic associations but is unlikely to create false positives.

### Genome scan of genomic regions associated with phenotypic differentiation in European local pig breeds

The genome scans for selection associated to broad phenotypic groups revealed 234 regions associated with stature, fatness, growth or reproduction traits. However, the distribution of the number of regions across traits is highly imbalanced, the reproduction phenotypic group containing 192 regions, while the growth traits only two. While these numbers could change slightly depending on the definition of a “significant region”, it is clear that the reproduction group exhibits more significant regions than the others. We could explain this result by different, non-exclusive reasons. First, it seems that our permutation procedure to correct p-values is less efficient for this breed score than for the others. This can be seen in the slightly biased distribution of p-values on the pp-plot of Figure 5 (Panel D-right). But another possibility is that the reproductive traits are more influenced by the production systems than the other traits. Therefore, testing for an association with the production system actually captures complex adaptations to other, non-measured traits than those linked to reproduction. The consequence is that by using this reproductive score, we are actually capturing adaption to many other traits linked to adaptation to the rearing conditions. Therefore, a bigger portion of the genome can be contributing to the response of local breeds to selection which could explain why p-values are generally smaller for this trait group.

Relatively little overlap was found between the significant regions discovered with the trait-association approach here and the trait agnostic one applied to the same pool sequencing data (8). Only two regions were found clearly overlapping, one on chromosome 8, encompassing the *MAP9* gene but also close to the *KIT* gene, associated notably to coat color, and the other on chromosome 15, containing genes *TMEM237* and *MPP4*. This illustrates the complementarity of the two approaches, and highlights empirically the gain in power that can be obtained by testing specific hypotheses (*e.g.* the association of allele frequencies to phenotypic variation) as was shown in simulations studies (20,21).

A strong association was found for stature and fatness in a region on chromosome 3 containing the *ANXA4* gene. Annexin A4 encodes a calcium-dependent phospholipid-binding protein involved in various membrane processes. This candidate gene or region has been previously proposed as a QTL for stature in cattle (46,47). Another genomic region associated with the signature of selection for stature and growth was discovered on chromosome 3 containing also the candidate gene *ANTXR1*. This gene is involved in cell morphogenesis, cellular development process and cytoskeleton organisation. Part of this region has also been previously proposed as a QTL for bovine stature (48). In the fatness and also stature group, two overlapping regions were found on chromosome 3. These regions contained common candidate genes; namely *DNMT3A* (responsible for CpG methylation), *POMC* (prohormone) and *EFR3B* (localises phosphatidylinositol 4-kinase to the plasma membrane). *DNMT3A* has been previously shown to affect stature and body weight in cattle and humans (48–50). In addition, the *DNMT3A* gene has previously been implicated to the regulation of adipose tissue development (51). Another candidate gene found in the same region is the *POMC* gene, which encodes the precursor of several peptide hormones that contribute to the regulation of feed intake and energy balance via the leptin/melanocortin pathway (52). Polymorphisms in *POMC* have previously been associated with *longissimus dorsi* muscle area and backfat thickness in cattle (53–55) and with obesity and body mass index in humans (56,57). An interesting signature of selection on reproductive performance was found on chromosome 4, which contains the *HSD17B7* gene. This gene encodes an enzyme involved in the biosynthesis of sex steroids and cholesterol (58,59). Due to the clear role of the *HSD17B7* gene in steroidogenesis, it is a good candidate gene for reproductive performance in local pig breeds.

Several other regions have also been identified as selection signatures for stature and fatness with not so direct connection to their biological function in production traits, but still with some association to phenotype. These regions could be useful for further/future studies on selection signatures. For example, chromosome 6 contained a region with possible adaptation on stature with the gene *ARHGAP28*, which has previously been associated with the number of vertebrae in pigs, thus affecting carcass length (60). In the fatness group, the gene *EIF2AK1* (located on chromosome 3) was identified, with a role of inhibiting of the protein synthesis in response to stress. This gene has previously been associated to body mass index in pigs (61).

Some of the regions that have been discovered must be carefully interpreted. For instance, the conducted studies differ in production conditions. Therefore, correlations between phenotypic groups could create non-causal signal with genes. An example of a non-causal signal due to phenotypic correlation found in the growth and fatness groups is a region on chromosome 8 containing the gene *KIT*. This gene encodes the tyrosine kinase receptor and is associated with coat colour in pigs. Another selection signature associated with coat colour was found on chromosome chromosome 6 in the growth and fatness groups. It contains the *MC1R* gene, which plays a major role in controlling the transition from eumelanin (black or brown) to pheomelanin (yellow to red) (62). Since the coat colour was favoured by breeders, it was strongly selected in several local breeds (63) (e.g. White breeds were under selection for leanness and better growth performance). Interestingly, the study performed on SNP-chip data on the same animals/breeds (9) did not detect any signal near *MC1R* or *KIT* genes probably due to different SNP-chip informativity and statistical approach used in their study.

### Conclusions

In this study, DNA-pool sequencing data of European local pig breeds were used to identify genomic regions associated with phenotypic differences in groups of traits related to stature, growth, fatness and reproductive performance. The genome scan for selection revealed several strong candidate genes with potential implication in adaptation of European local pig breeds to different production systems. This study will be helpful for future conservation, association or selection approaches in these European local pig breeds.

## Supporting information

Supplementary Figurtes

PCA for all breeds

## Declarations

### Ethics approval and consent to participate

Not applicable.

### Consent for publication

Not applicable.

### Availability of data and materials

Data used in this study are available from the original sources as detailed in the Materials and Methods section.

## Authors’ contributions

BS, MČP and MŠ conceived the study. MČP and MŠ obtained project funding and were responsible for project administration. BS supervised the study and designed the methodology. CO, LF, SB, GS, AR, MM, MG, RB, RC, RQ, GK, MJM, CZ, VR, JPA, ČR, RS, DK provided data and samples. KP, CM, BS performed data analysis. KP, CM, BS, MČP, MŠ, LF, CO and JR contributed to data interpretation. KP, CM and BS wrote the initial draft of the paper. All authors reviewed and edited paper. All authors read and approved the final manuscript.

## Acknowledgements

This study has received funding from the European Union’s Horizon 2020 research and innovation programme under grant agreement No 634476 (project acronym TREASURE). Project TREASURE consortium is acknowledged for providing the samples. The content of this paper reflects only the author’s view and the European Union Agency is not responsible for any use that may be made of the information it contains. Core financing of the Slovenian Research Agency (grant P4-0133, J4-3094, PhD scholarship for KP) for KP, MČP and MŠ is acknowledged. The authors also wish to thank Nina Batorek-Lukač and Urška Tomažin for their contribution in phenotypic data collection.

## Additional files

### Additional file 1 Table S1

Title: Summary of sequencing statistics (adapted from Bovo et al., 2020).

### Additional file 1 Table S2

Title: Description of phenotypic traits used for phenotypic characterisation of European local pig breeds.

### Additional file 2 Figure S1

Title: Plot od admixture cross-validation error from K=2 to K=40.

### Additional file 2 Figure S2

Title: Admixture plots of European local pig breeds for K=24, K=20, K=15, K=10, and K=6 (from the outside to the inside of the circle).

### Additional file 2 Figure S3

Title: Projection of the pool-sequencing data (allele frequencies) on the PCA established with individual genotypes on the SNP genotyping array. The x-axis is the average coordinate of individuals of each population on each of the PC of the genotyped-based PCA, and the y-axis is the coordinate of the allele frequencies of each population projected on each PCA.

### Additional file 2 Figure S4

Title: A matrix of missing phenotypic variables in a database of European local pig breeds. The yellow colour is representing missing variables.

Description: SFA = saturated fatty acid content, PUFA = polyunsaturated fatty acid content, Piglets W weight = piglets weaning weight, MUFA = monounsaturated fatty acid content, Litter W birth = litter weaning weight, LD IMF = *longissimus dorsi* intramuscular fat content, M = male, F = female, Death WN = death rate to weaning, BFT = backfat thickness, GM = *gluteus medius* muscle, ADG1 = Average daily gain during the lactation period, ADG2 = Average daily gain in the growing period from weaning to 30 kg of weight, ADG3 = Average daily gain during fattening period from 30 kg to 60 kg.

### Additional file 2 Figure S5

Title: Principal component analysis showing the relationship between breeds (A) and the traits associated with growth (B) and the corresponding phenotypic breed scores (C).

Description: Breeds (A) coloured in grey are the breeds with more than 50% of missing variables, thus, their position on the PCA must be interpreted carefully. The variables (B) are coloured according to quality of the representation, which is measured by squared cosine between the vector originating from the element and its projection on the axis. The variables that contribute most to the separation of the trait into PC1 and PC2 are coloured black. Breeds (A, C) are coloured according to genetic similarity. Breeds (A, C) in green are genetically Landrace-like breeds, in purple are Large White-like breeds, in blue are Iberian-like breeds, in red are Duroc-like breeds and in light blue are Gascon and Basque.

### Additional file 2 Figure S6

Title: Principal component analysis showing the relationship between breeds (A) and the traits associated with fatness (B) and the corresponding phenotypic breed scores (C).

Description: Breeds (A) coloured in grey are the breeds with more than 50% of missing variables, thus, their position on the PCA must be interpreted carefully. The variables (B) are coloured according to quality of the representation, which is measured by squared cosine between the vector originating from the element and its projection on the axis. The variables that contribute most to the separation of the trait into PC1 and PC2 are coloured black. Breeds (A, C) are coloured according to genetic similarity. Breeds (A, C) in green are genetically Landrace-like breeds, in purple are Large White-like breeds, in blue are Iberian-like breeds, in red are Duroc-like breeds and in light blue are Gascon and Basque.

### Additional file 2 Figure S7

Title: Principal component analysis showing the relationship between breeds (A) and the traits associated with reproduction performance (B) and the corresponding phenotypic breed scores (C).

Description: Breeds (A) coloured in grey are the breeds with more than 50% of missing variables, thus, their position on the PCA must be interpreted carefully. The variables (B) are coloured according to quality of the representation, which is measured by squared cosine between the vector originating from the element and its projection on the axis. The variables that contribute most to the separation of the trait into PC1 and PC2 are coloured black. Breeds (A, C) are coloured according to genetic similarity. Breeds (A, C) in green are genetically Landrace-like breeds, in purple are Large White-like breeds, in blue are Iberian-like breeds, in red are Duroc-like breeds and in light blue are Gascon and Basque.

### Additional file 2 Figure S8

Title: Distribution of the European local pig breeds according to intramuscular fat content in longissimus dorsi (LD) muscle.

Description: Higher values on x-axis are representing higher intramuscular fat content, while the lower values are representing lower intramuscular fat content.

### Additional file 2 Figure S9

Title: Circular plot summarizing regions with candidate genes for phenotypic groups growth, stature, fatness and reproduction.

Description: Discovered regions associated with growth are coloured in red, with stature are coloured in blue, with fatness are coloured in purple and with reproductive performance in green.

### Additional file 2 Figure S10

Title: Principal component analysis showing the relationship between breeds (A) and the traits associated with meat quality (B) and the corresponding phenotypic breed scores (C).

Description: Breeds (A) coloured in grey are the breeds with more than 50% of missing variables, thus, their position on the PCA must be interpreted carefully. The variables (B) are coloured according to quality of the representation, which is measured by squared cosine between the vector originating from the element and its projection on the axis. The variables that contribute most to the separation of the trait into PC1 and PC2 are coloured black. Breeds (A, C) are coloured according to genetic similarity. Breeds (A, C) in green are genetically Landrace-like breeds, in purple are Large White-like breeds, in blue are Iberian-like breeds, in red are Duroc-like breeds and in light blue are Gascon and Basque.

### Additional file 3

Title: The genetic structure of European local pig populations assessed from individual SNP genotyping data using principal component analysis.

### Additional file 4

Title: A list of detected genomic regions associated with phenotypic differentiation in production traits in European local pig breeds.

Format: xls

