## Supplementary Tables for "Discovering genomic regions associated with the phenotypic differentiation of European local pig breeds"

Additional information 1

Additional File 1: Table S1

Additional File 1: Table S2

**Table S1.** Summary of sequencing statistics (adapted from Bovo et. al, 2020)

| Population | No. of read pairs | Depth of coverage * |
|---|---|---|
| Alentejano | 419,690,476 | 41.98 |
| Apulo-Calabrese | 418,529,727 | 42.12 |
| Basque | 407,698,128 | 39.55 |
| Bísaro | 415,284,437 | 42.44 |
| Black Slavonian | 405,316,112 | 40.61 |
| Casertana | 435,598,516 | 43.61 |
| Cinta Senese | 422,120,850 | 42.42 |
| Gascon | 408,764,207 | 41.10 |
| Krškopolje | 404,204,144 | 40.80 |
| Lietuvus Vietiné | 409,935,460 | 41.99 |
| Lietuvus Baltosios Senojo Tipo | 405,822,217 | 41.62 |
| Negre Mallorquí | 414,314,159 | 41.92 |
| Mora Romagnola | 411,095,541 | 41.21 |
| Moravka | 413,100,992 | 42.27 |
| Nero Siciliano | 405,812,223 | 38.92 |
| Sarda | 442,035,147 | 44.32 |
| Schwäbisch-Hällisches | 428,982,876 | 42.69 |
| Mangalitsa | 416,663,891 | 41.08 |
| Turopolje | 416,663,891 | 42.61 |
| Italian Duroc | 420,384,723 | 41.91 |
| Italian Landrace | 442,780,637 | 44.35 |
| Italian Large White | 450,673,024 | 45.24 |

* after removal of duplicated reads

**Table S2.** Description of phenotypic traits used for phenotypic characterisation of European local pig breeds.

| Group | Trait code | Description of the trait |
|---|---|---|
| Growth performance | ADG 1 | Average daily gain during the lactation period. |
| | ADG 2 | Growing period from weaning to 30 kg of weight. |
| | ADG 3 | First fattening period from 30 to 60 kg, |
| Stature | Bodyweight - M | Average male body weight. |
| | Bodyweight - F | Average female body weight. |
| | Height at withers - M | Male height at withers. |
| | Height at withers -F | Female height at withers. |
| Fatness | BFT last rib | Backfat thickness at the level of the last rib. |
| | BFT withers | Backfat thickness on withers. |
| | BFT at GM | Backfat thickness above *gluteus medius*. |
| | LD IMF | Intramuscular fat content in *longissimus dorsi* muscle. |
| | Meatiness | Meat content. |
| | Loin eye area | Loin eye area. |
| | SFA | Saturated fatty acid content of *longissimus dorsi* muscle. |
| | MUFA | Monounsaturated fatty acid content of *longissimus dorsi* muscle. |
| | PUFA | Polyunsaturated fatty acid content of *longissimus dorsi* muscle. |
| | BFT last rib LM | Adjusted values for backfat thickness at the level of the last rib on final body weight of 120 kg using linear mixed models. |
| Reproductive performance | Sow age | Age of sow in months at first parturition. |
| | Litter/year | Litters per sow per year. |
| | L W birth | Litter weight at birth in kg. |
| | Piglets/litter | Piglets per litter. |
| | Alive piglets/litter | Piglets alive per litter. |
| | Piglet live BW | Piglet live birth weight in kg. |
| | Stillborn/litter | Stillborn pigs per litter. |
| | Death WN | Death rate to weaning. |
| | Weaned piglets/litter | Piglets weaned per litter. |
| | Piglets W weight | Piglets weaning weight. |
| | Lactation | Duration of lactation. |
| | Farrowing interval | Farrowing interval in days. |
