## Supplementary Figurtes for "Discovering genomic regions associated with the phenotypic differentiation of European local pig breeds"

Additional information 2

Additional File 2: Figure S1

Additional File 2: Figure S2

Additional File 2: Figure S3

Additional File 2: Figure S4

Additional File 2: Figure S5

Additional File 2: Figure S6

Additional File 2: Figure S7

Additional File 2: Figure S8

Additional File 2: Figure S9

Additional File 2: Figure S10

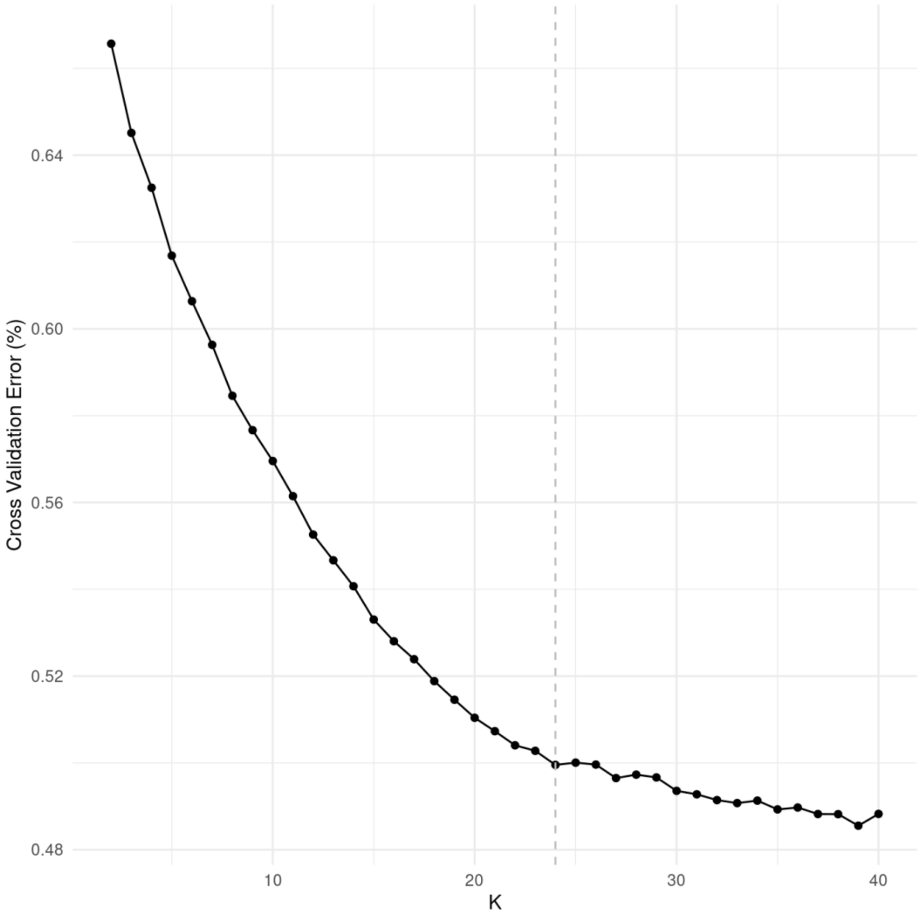


**Figure S1.** Plot of admixture cross-validation error from K=2 to K=40.

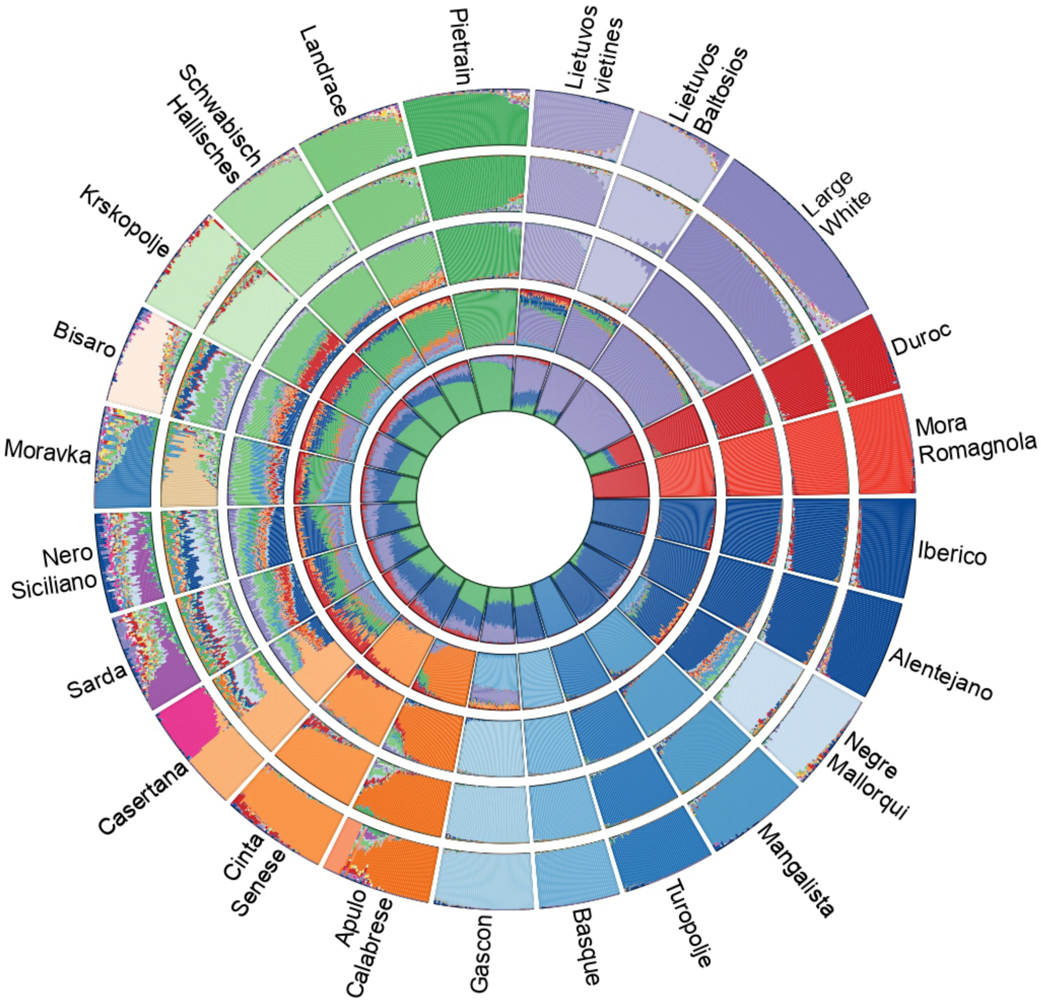


**Figure S2:** Admixture plots of European local pig breeds for K=24, K=20, K=15, K=10, and K=6 (from the outside to the inside of the circle).

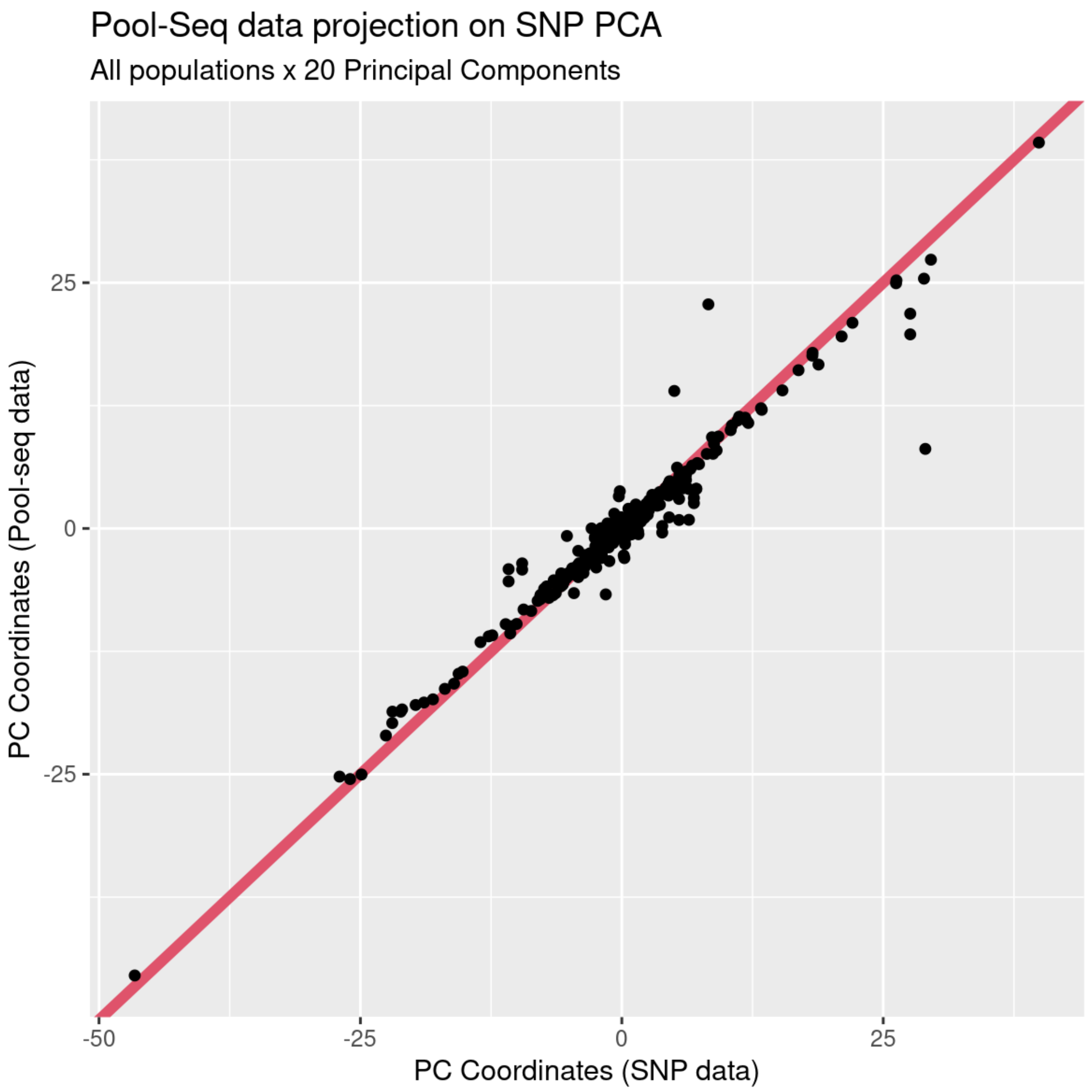


**Figure S3:** Projection of the pool-sequencing data (allele frequencies) on the PCA established with individual genotypes on the SNP genotyping array. The x-axis is the average coordinate of individuals of each population on each of the PC of the genotyped-based PCA, and the y-axis is the coordinate of the allele frequencies of each population projected on each PCA.

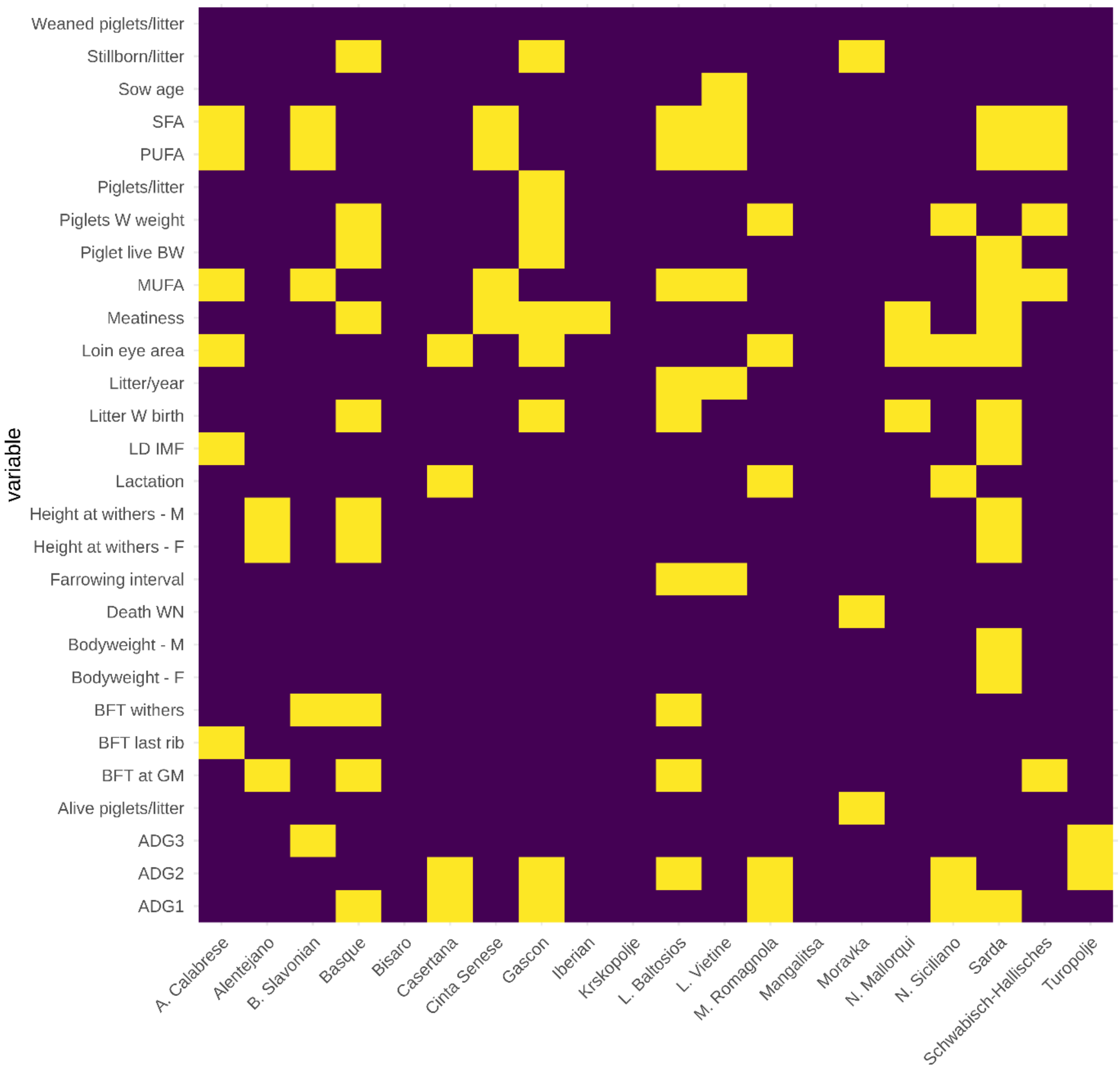


**Figure S4.** A matrix of missing phenotypic variables in a database of European local pig breeds. The yellow colour is representing missing variables. SFA = saturated fatty acid content, PUFA = polyunsaturated fatty acid content, Piglets W weight = piglets weaning weight, MUFA = monounsaturated fatty acid content, Litter W birth = litter weaning weight, LD IMF = *longissimus dorsi* intramuscular fat content, M = male, F = female, Death WN = death rate to weaning, BFT = backfat thickness, GM = *gluteus medius* muscle, ADG1 = Average daily gain during the lactation period, ADG2 = Average daily gain in the growing period from weaning to 30 kg of weight, ADG3 = Average daily gain during fattening period from 30 kg to 60 kg.

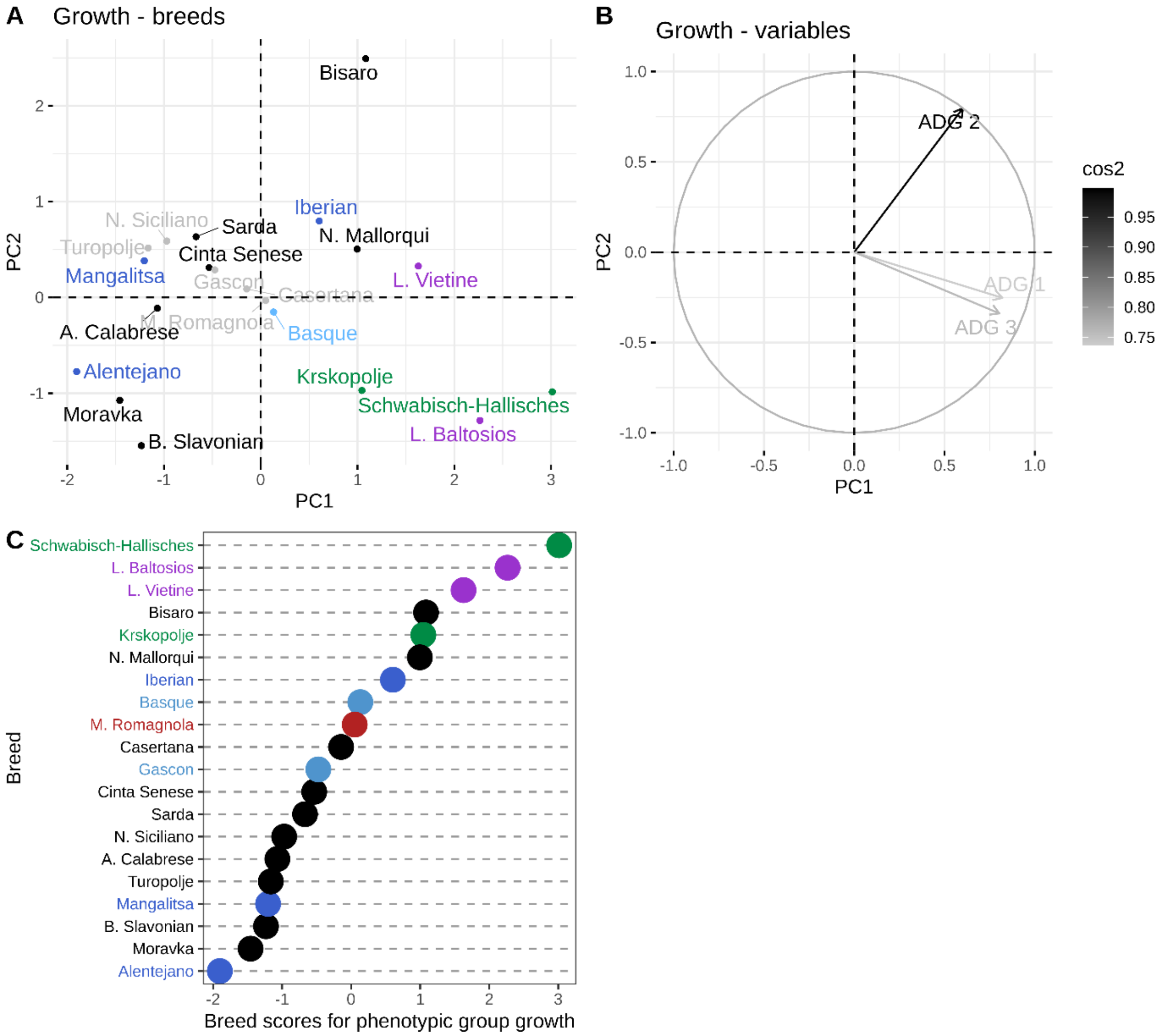


**Figure S5.** Principal component analysis showing the relationship between breeds (A) and the traits associated with growth (B) and the corresponding phenotypic breed scores (C).

Breeds (A) coloured in grey are the breeds with more than 50% of missing variables, thus, their position on the PCA must be interpreted carefully. The variables (B) are coloured according to quality of the representation, which is measured by squared cosine between the vector originating from the element and its projection on the axis. The variables that contribute most to the separation of the trait into PC1 and PC2 are coloured black. Breeds (A, C) are coloured according to genetic similarity. Breeds (A, C) in green are genetically Landrace-like breeds, in purple are Large White-like breeds, in blue are Iberian-like breeds, in red are Duroc-like breeds and in light blue are Gascon and Basque.

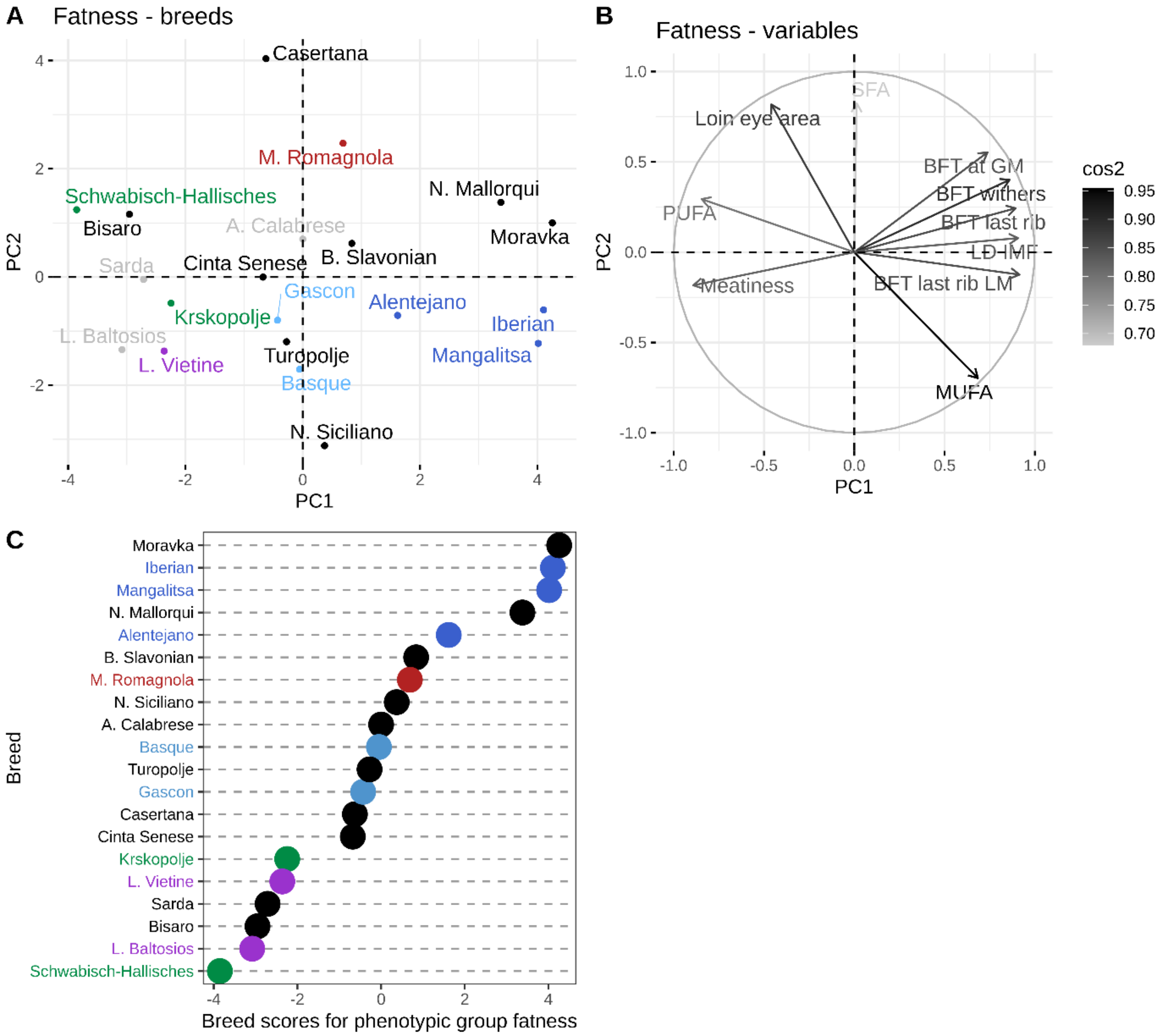


**Figure S6.** Principal component analysis showing the relationship between breeds (A) and the traits associated with fatness (B) and the corresponding phenotypic breed scores (C).

Breeds (A) coloured in grey are the breeds with more than 50% of missing variables, thus, their position on the PCA must be interpreted carefully. The variables (B) are coloured according to quality of the representation, which is measured by squared cosine between the vector originating from the element and its projection on the axis. The variables that contribute most to the separation of the trait into PC1 and PC2 are coloured black. Breeds (A, C) are coloured according to genetic similarity. Breeds (A, C) in green are genetically Landrace-like breeds, in purple are Large White-like breeds, in blue are Iberian-like breeds, in red are Duroc-like breeds and in light blue are Gascon and Basque.

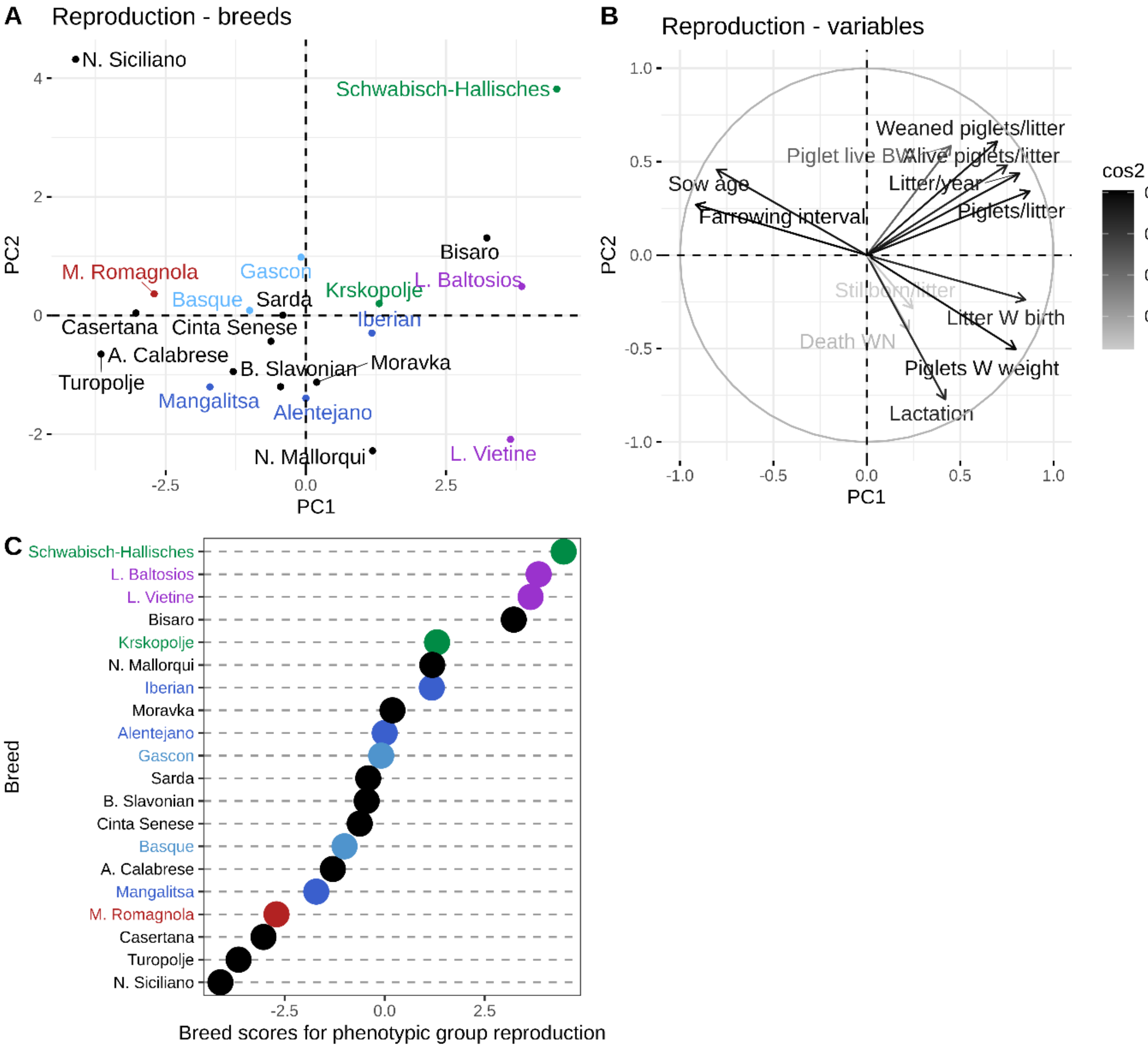


**Figure S7.** Principal component analysis showing the relationship between breeds (A) and the traits associated with reproduction performance (B) and the corresponding phenotypic breed scores (C).

Breeds (A) coloured in grey are the breeds with more than 50% of missing variables, thus, their position on the PCA must be interpreted carefully. The variables (B) are coloured according to quality of the representation, which is measured by squared cosine between the vector originating from the element and its projection on the axis. The variables that contribute most to the separation of the trait into PC1 and PC2 are coloured black. Breeds (A, C) are coloured according to genetic similarity. Breeds (A, C) in green are genetically Landrace-like breeds, in purple are Large White-like breeds, in blue are Iberian-like breeds, in red are Duroc-like breeds and in light blue are Gascon and Basque.

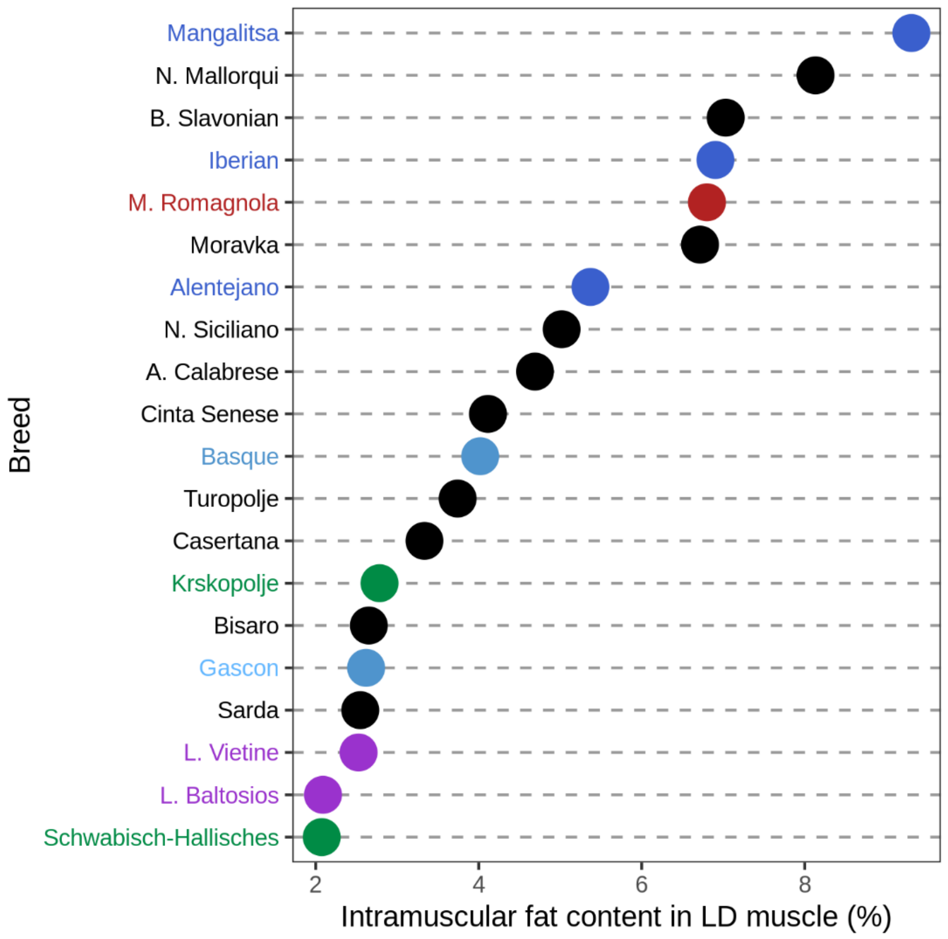


**Figure S8.** Distribution of the European local pig breeds according to intramuscular fat content in *longissimus dorsi* (LD) muscle. Higher values on x-axis are representing higher intramuscular fat content, while the lower values are representing lower intramuscular fat content.

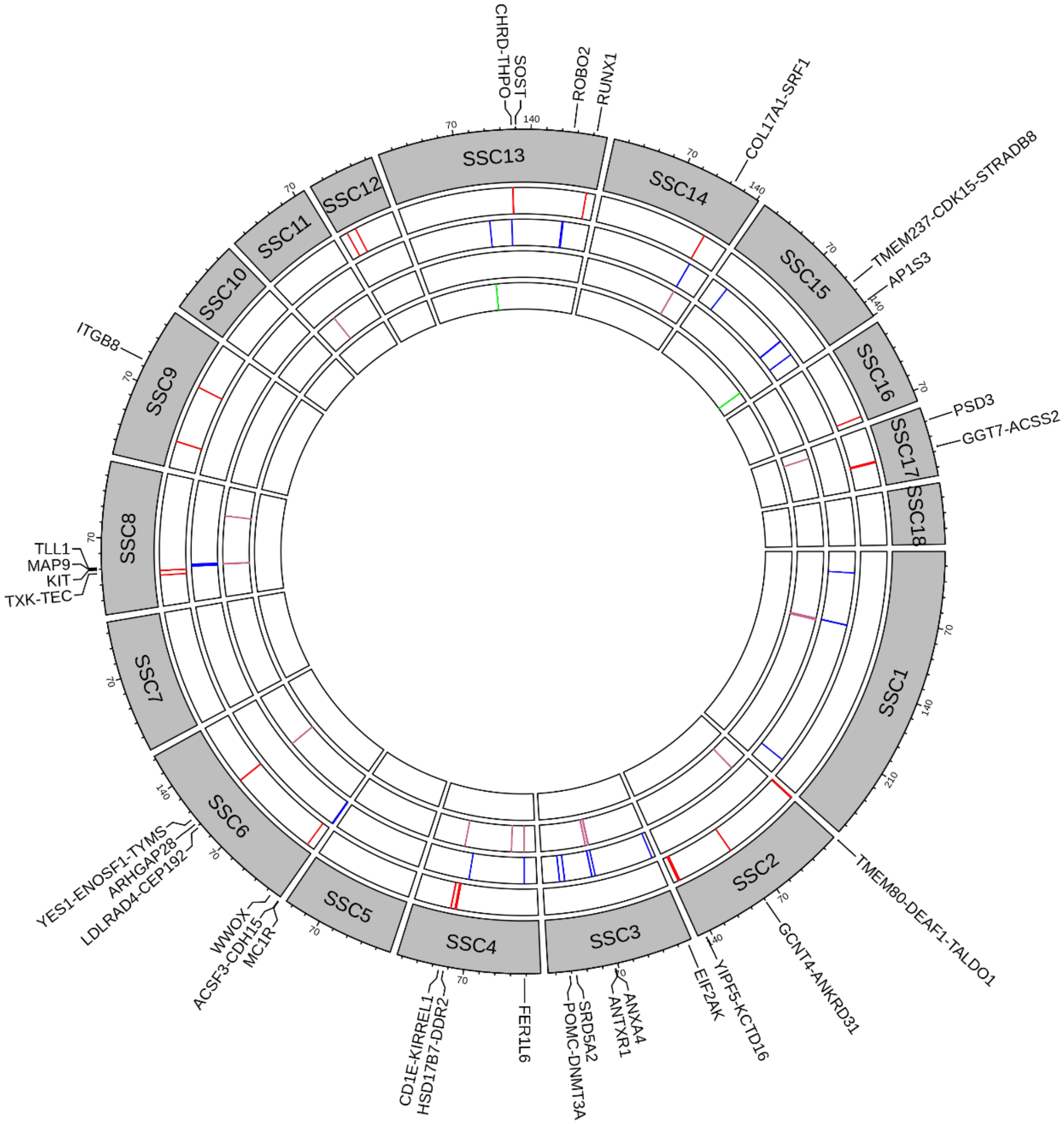


**Figure S9.** Circular plot summarizing regions with candidate genes for phenotypic groups growth, stature, fatness and reproduction. Discovered regions associated with growth are coloured in red, with stature are coloured in blue, with fatness are coloured in purple and with reproductive performance in green.

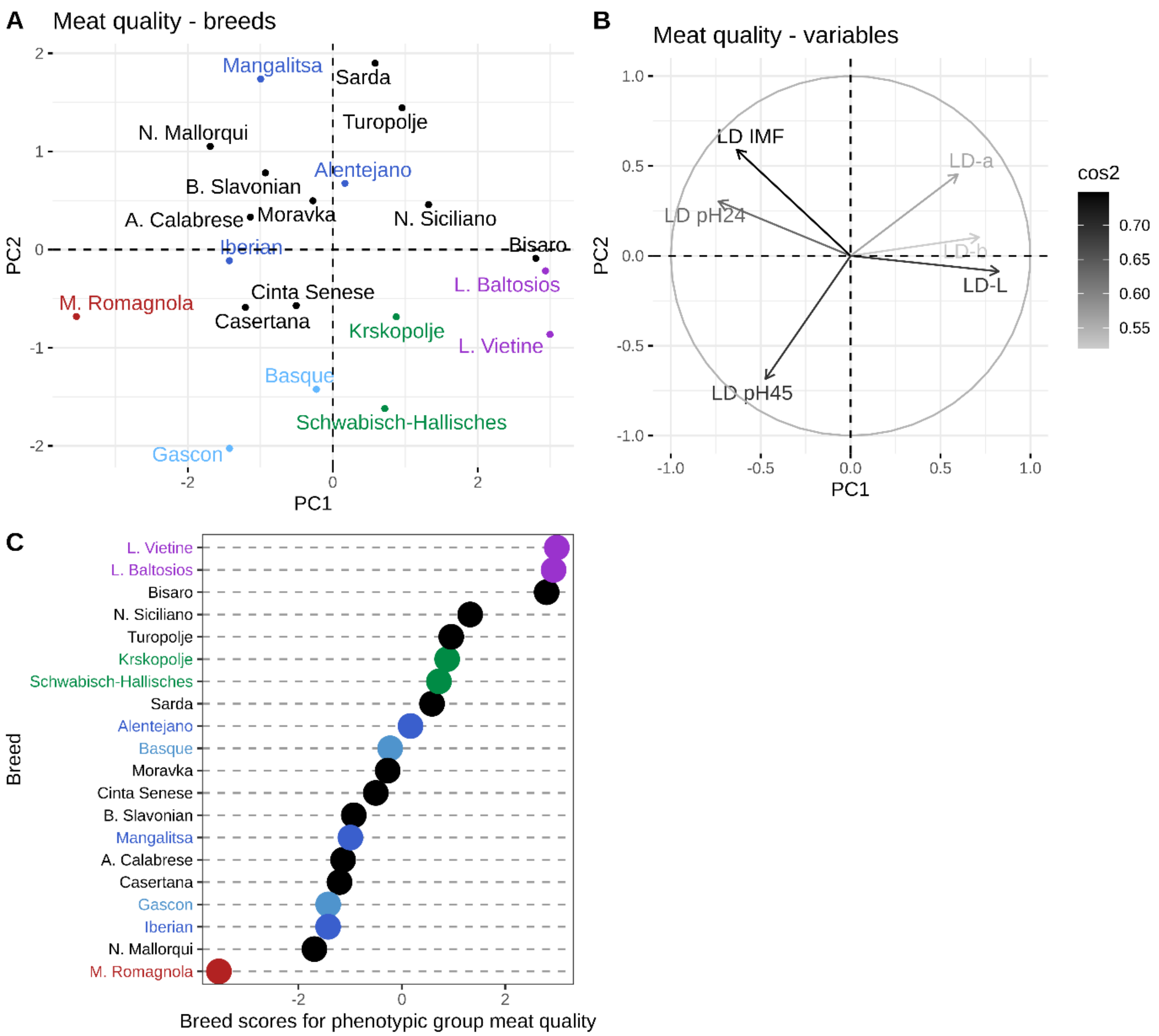


**Figure S10.** Principal component analysis showing the relationship between breeds (A) and the traits associated with meat quality (B) and the corresponding phenotypic breed scores (C). Breeds (A) coloured in grey are the breeds with more than 50% of missing variables, thus, their position on the PCA must be interpreted carefully. The variables (B) are coloured according to quality of the representation, which is measured by squared cosine between the vector originating from the element and its projection on the axis. The variables that contribute most to the separation of the trait into PC1 and PC2 are coloured black. Breeds (A, C) are coloured according to genetic similarity. Breeds (A, C) in green are genetically Landrace-like breeds, in purple are Large White-like breeds, in blue are Iberian-like breeds, in red are Duroc-like breeds and in light blue are Gascon and Basque.
