## Supplementary material for "Discovering genomic regions associated with the phenotypic differentiation of European local pig breeds": PCA for all breeds

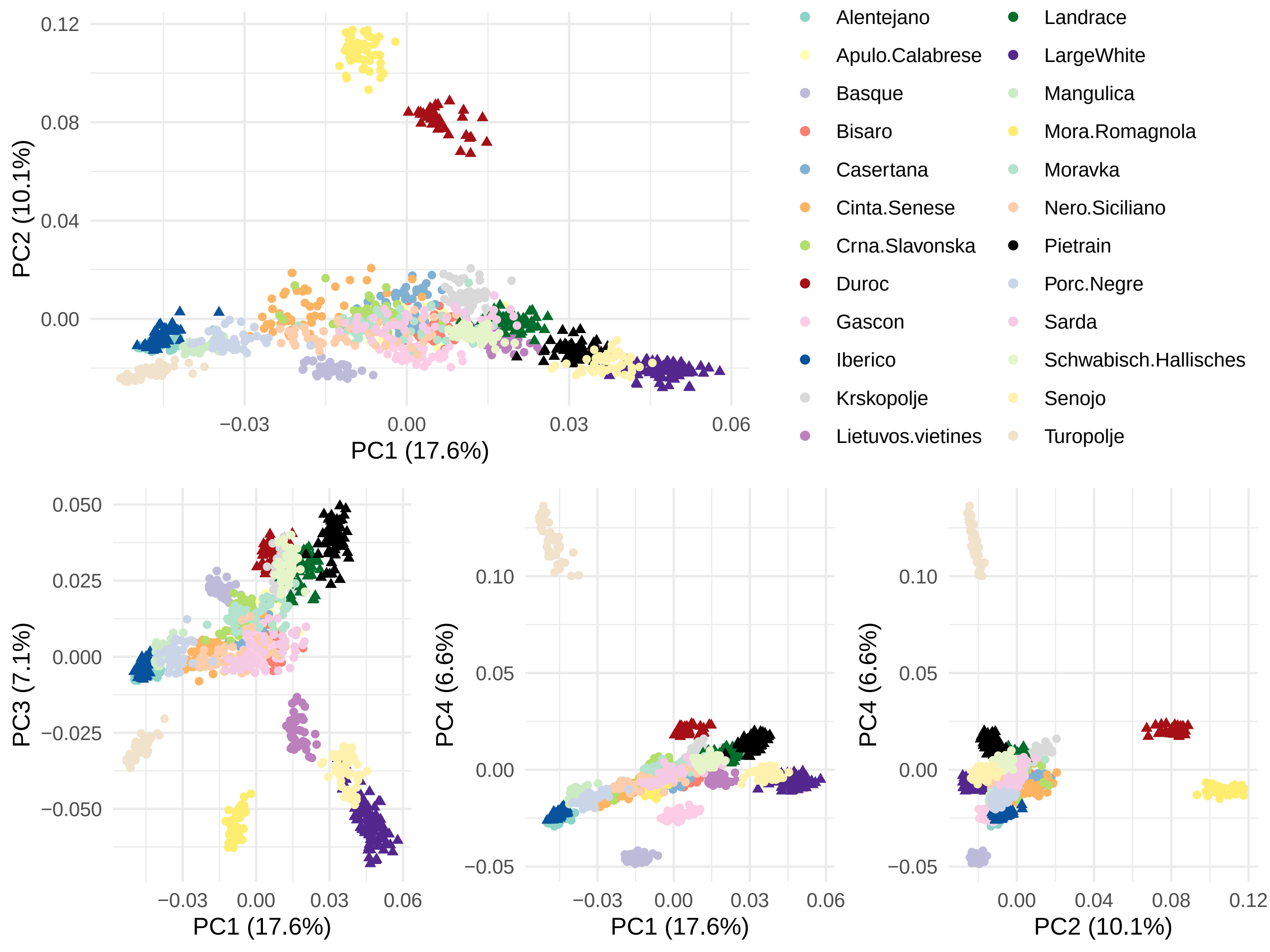

PC2 (10.1%)
PC1 (17.6%)
0.12
0.08
0.04
0.00
−0.03
0.00
0.03
0.06
Alentejano
Apulo.Calabrese
Basque
Bisaro
Casertana
Cinta.Senese
Crna.Slavonska
Duroc
Gascon
Iberico
Krskopolje
Lietuvos.vietines
Landrace
LargeWhite
Mangulica
Mora.Romagnola
Moravka
Nero.Siciliano
Pietrain
Porc.Negre
Sarda
Schwabisch.Hallisches
Senojo
Turopolje
PC3 (7.1%)
0.050
0.025
0.000
−0.025
−0.050
PC1 (17.6%)
PC4 (6.6%)
0.10
0.05
0.00
−0.05
PC1 (17.6%)
PC4 (6.6%)
PC2 (10.1%)
0.04
0.08
0.12

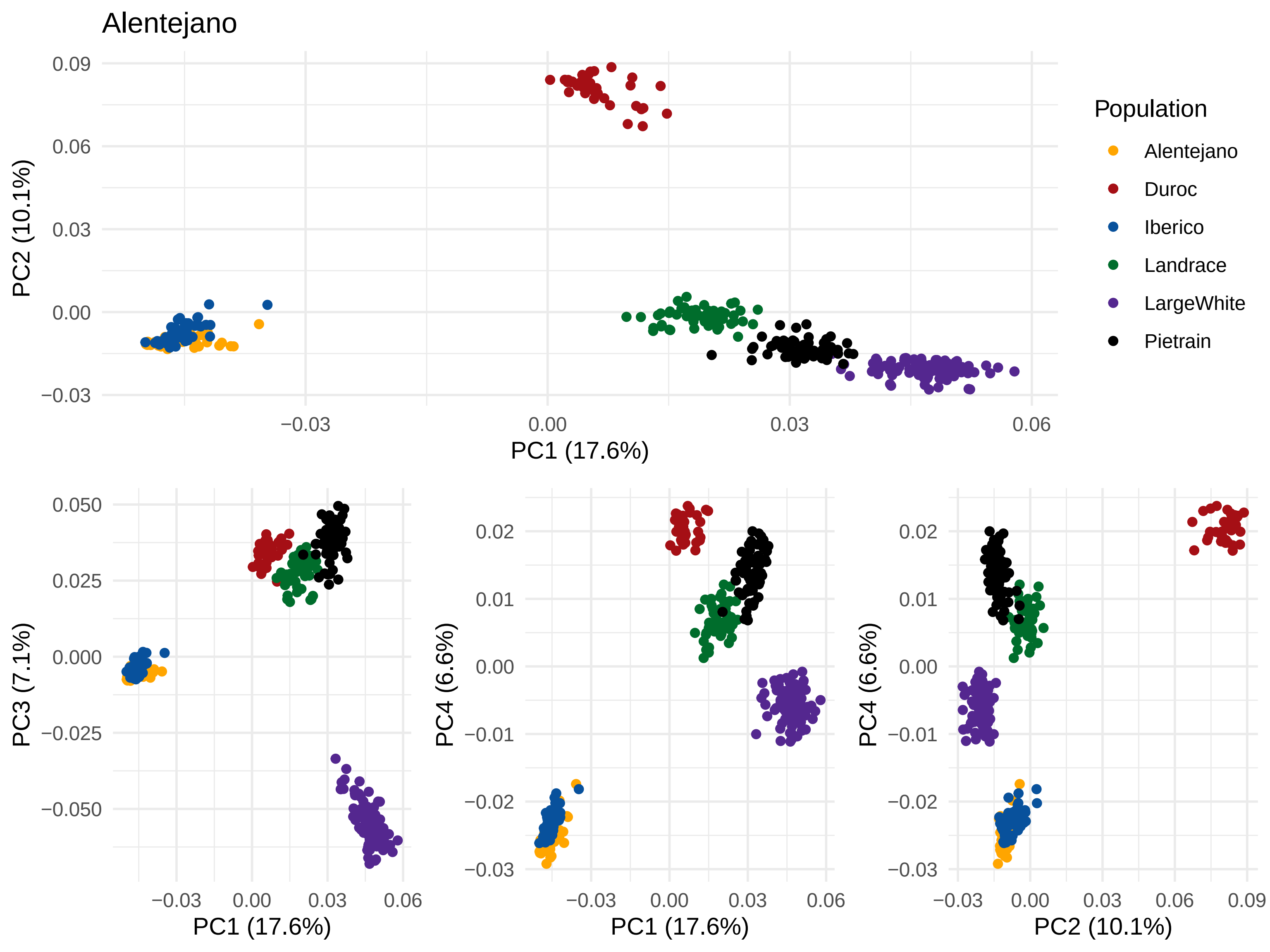

Alentejano
PC2 (10.1%)
PC1 (17.6%)
0.09
0.06
0.03
0.00
−0.03
0.00
0.03
0.06
Population
Alentejano
Duroc
Iberico
Landrace
LargeWhite
Pietrain
PC3 (7.1%)
0.050
0.025
0.000
−0.025
−0.050
PC4 (6.6%)
0.02
0.01
0.00
−0.01
−0.02
−0.03
PC2 (10.1%)
0.09

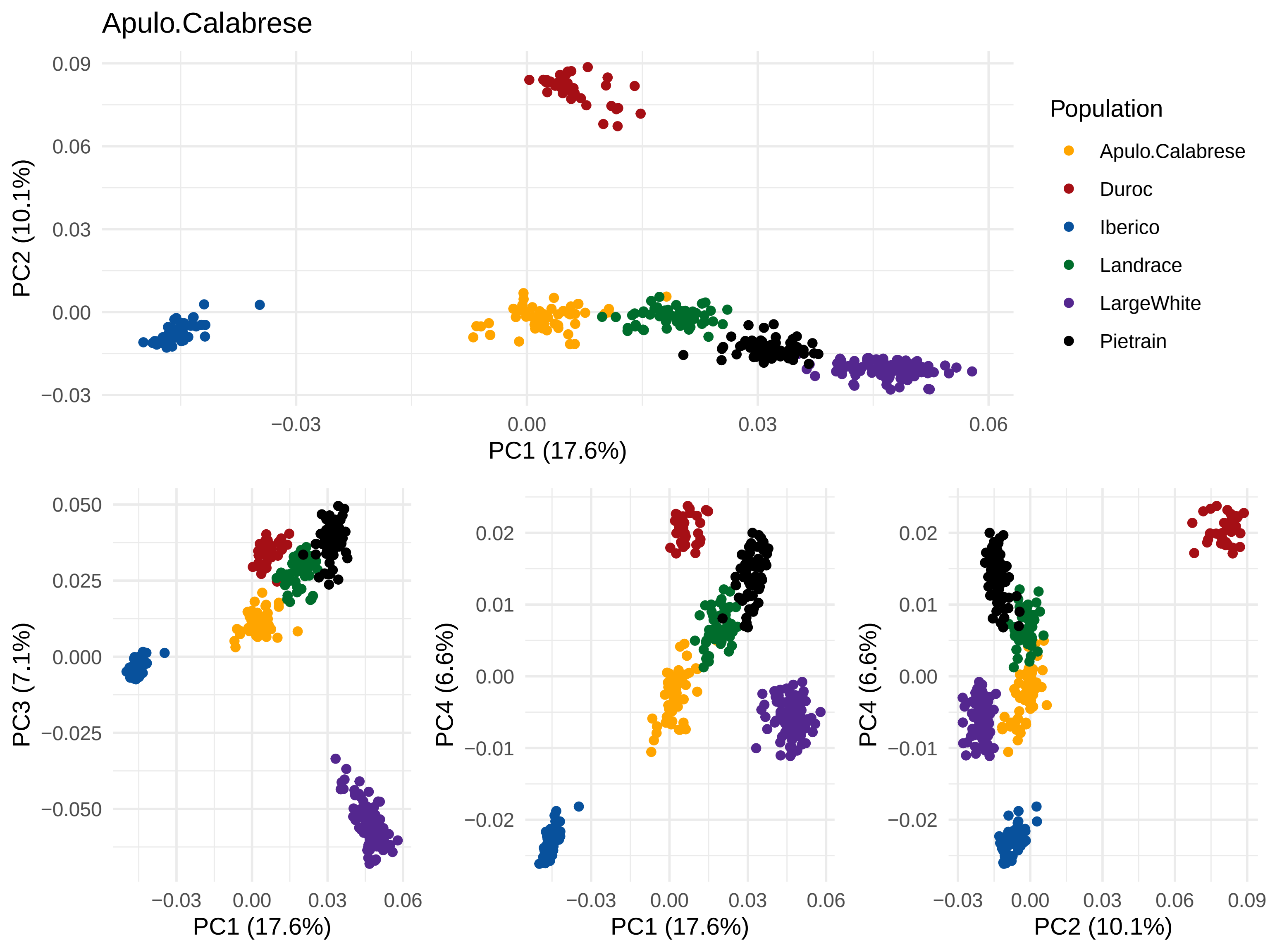
Apulo.Calabrese
PC2 (10.1%)
PC1 (17.6%)
Population
Apulo.Calabrese
Duroc
Iberico
Landrace
LargeWhite
Pietrain
PC3 (7.1%)
PC4 (6.6%)
PC1 (17.6%)
PC1 (17.6%)
PC4 (6.6%)
PC2 (10.1%)

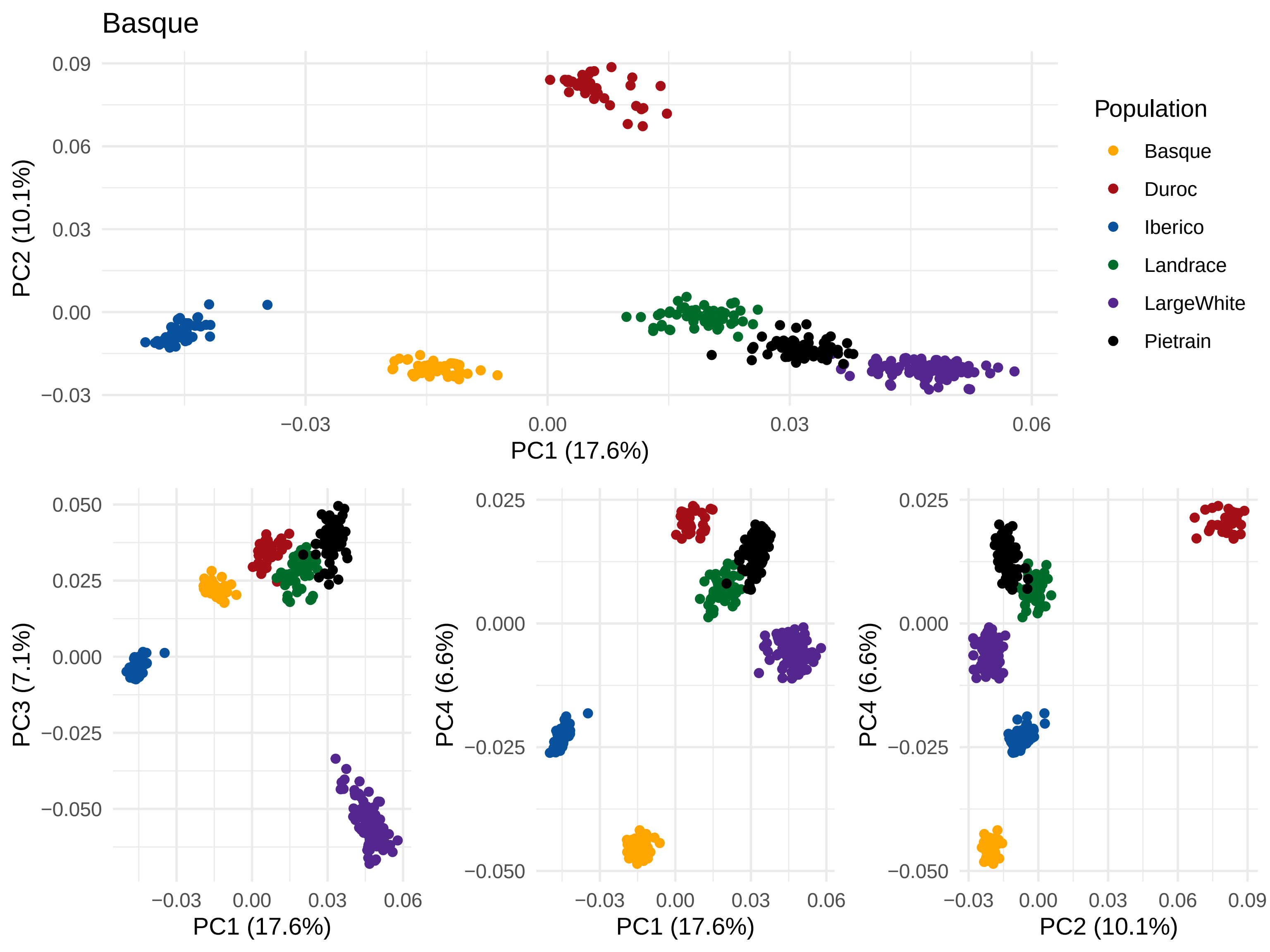
Basque
PC1 (17.6%)
PC2 (10.1%)
PC3 (7.1%)
PC4 (6.6%)
Population
Basque
Duroc
Iberico
Landrace
LargeWhite
Pietrain
0.09
0.06
0.03
0.00
−0.03
0.050
0.025
0.000
−0.025
−0.050
0.025
0.000
−0.025
−0.050
−0.03
0.00
0.03
0.06
0.09

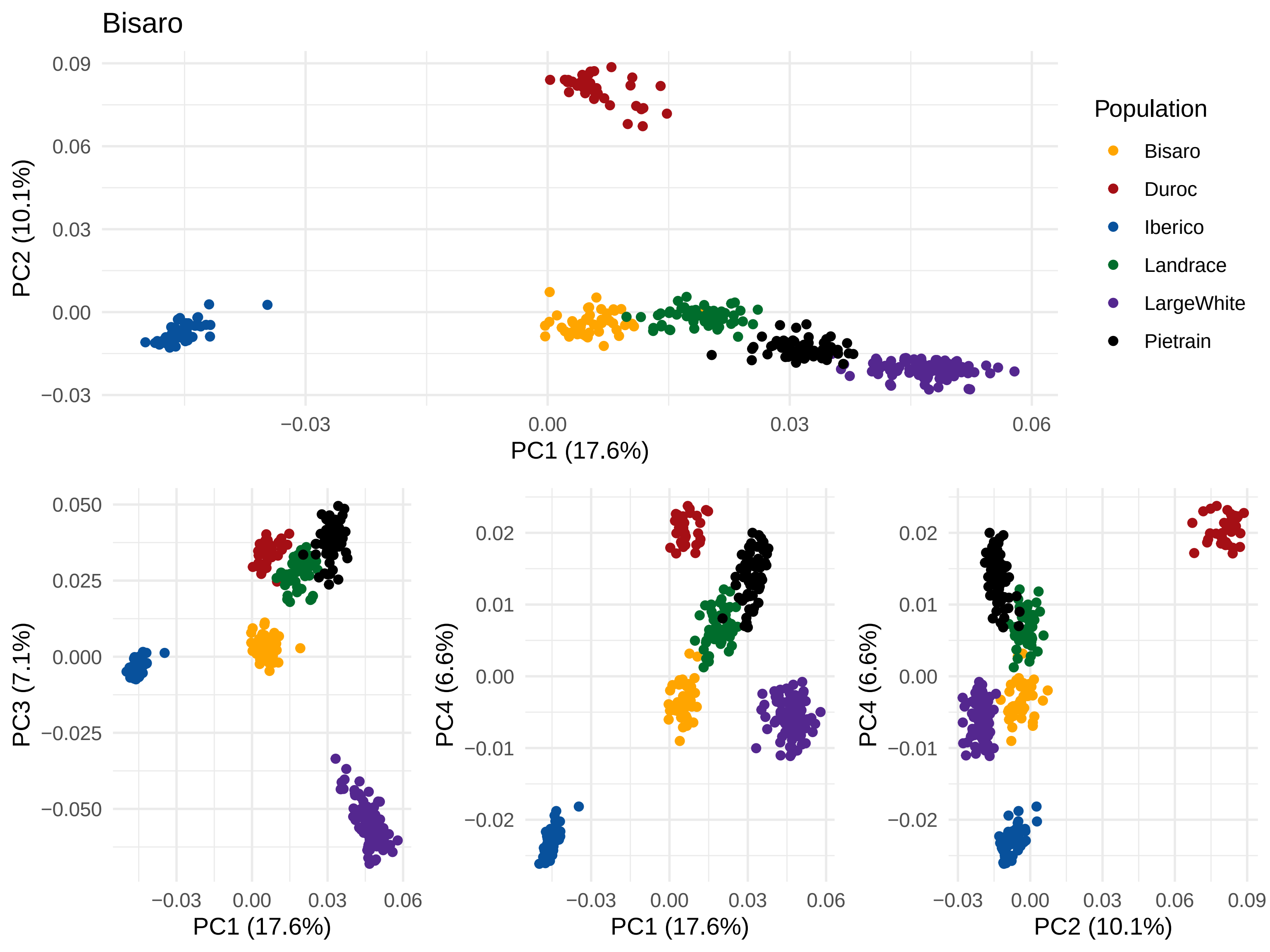
Bisaro
PC1 (17.6%)
PC2 (10.1%)
PC3 (7.1%)
PC4 (6.6%)
Population
Bisaro
Duroc
Iberico
Landrace
LargeWhite
Pietrain

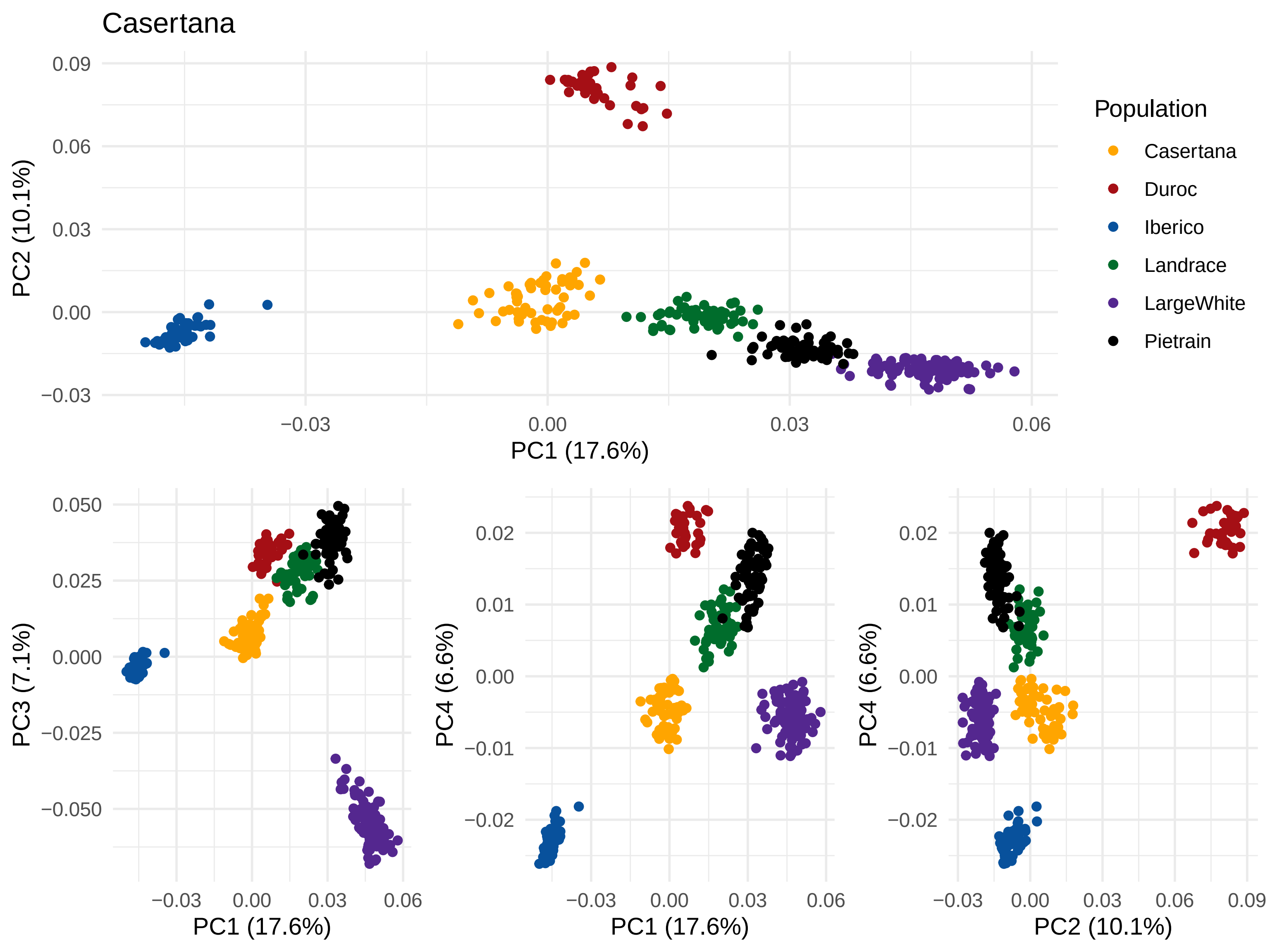

Casertana
PC2 (10.1%)
PC1 (17.6%)
Population
Casertana
Duroc
Iberico
Landrace
LargeWhite
Pietrain
PC3 (7.1%)
PC4 (6.6%)
PC1 (17.6%)
PC1 (17.6%)
PC4 (6.6%)
PC2 (10.1%)

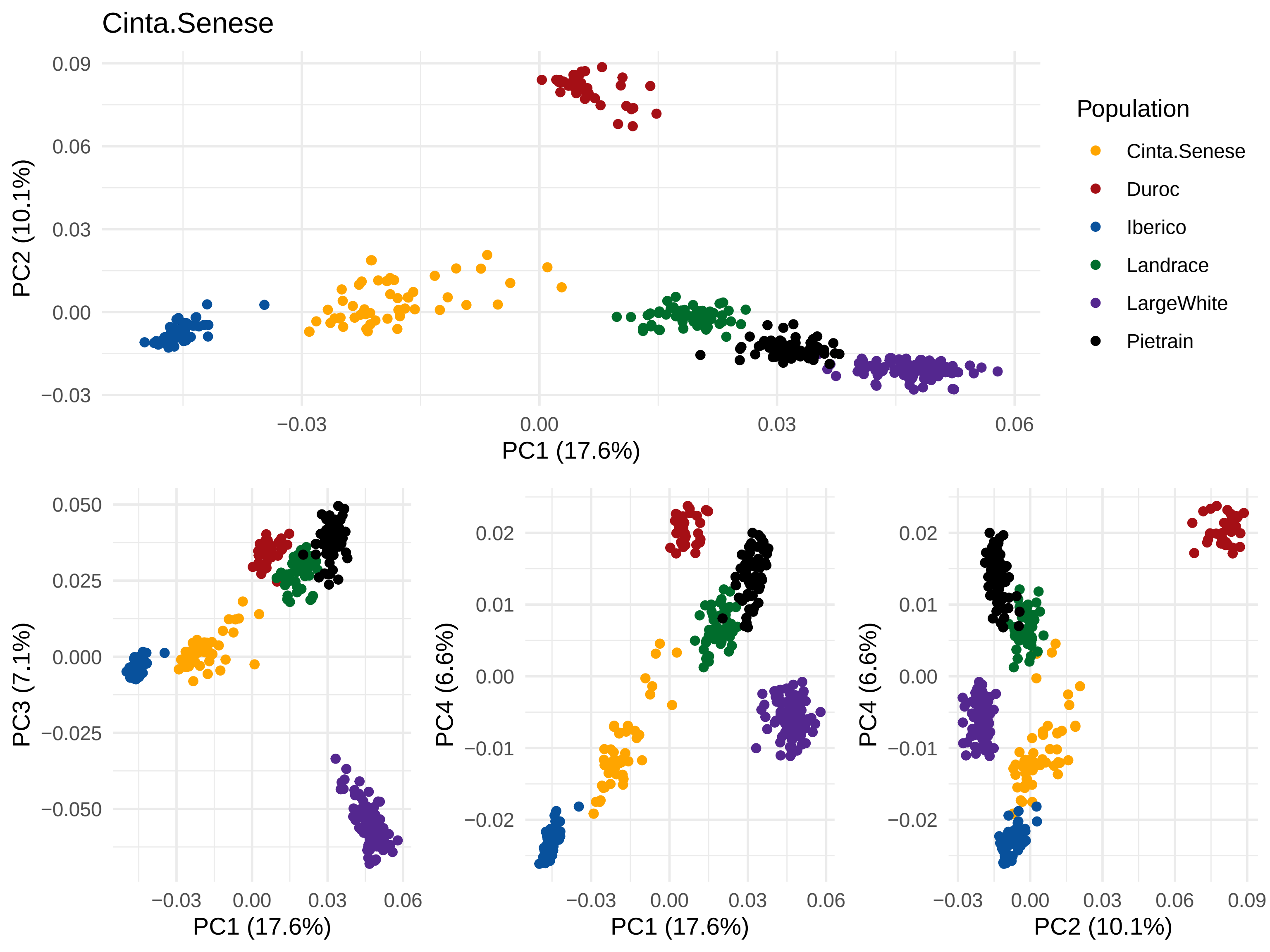

Cinta.Senese
PC1 (17.6%)
PC2 (10.1%)
PC3 (7.1%)
PC4 (6.6%)
Population
Cinta.Senese
Duroc
Iberico
Landrace
LargeWhite
Pietrain

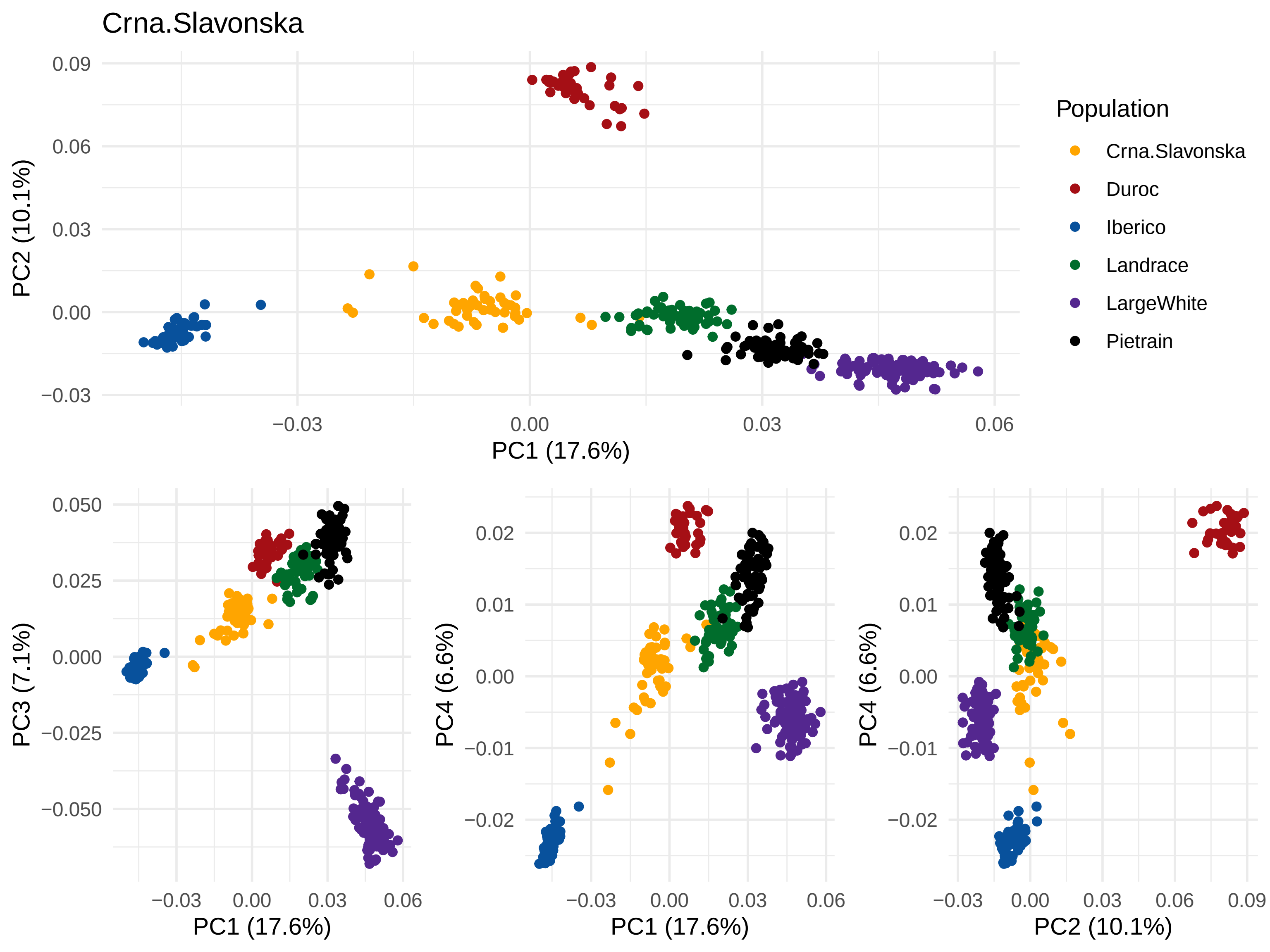
Crna.Slavonska
PC1 (17.6%)
PC2 (10.1%)
PC3 (7.1%)
PC4 (6.6%)
Population
Crna.Slavonska
Duroc
Iberico
Landrace
LargeWhite
Pietrain

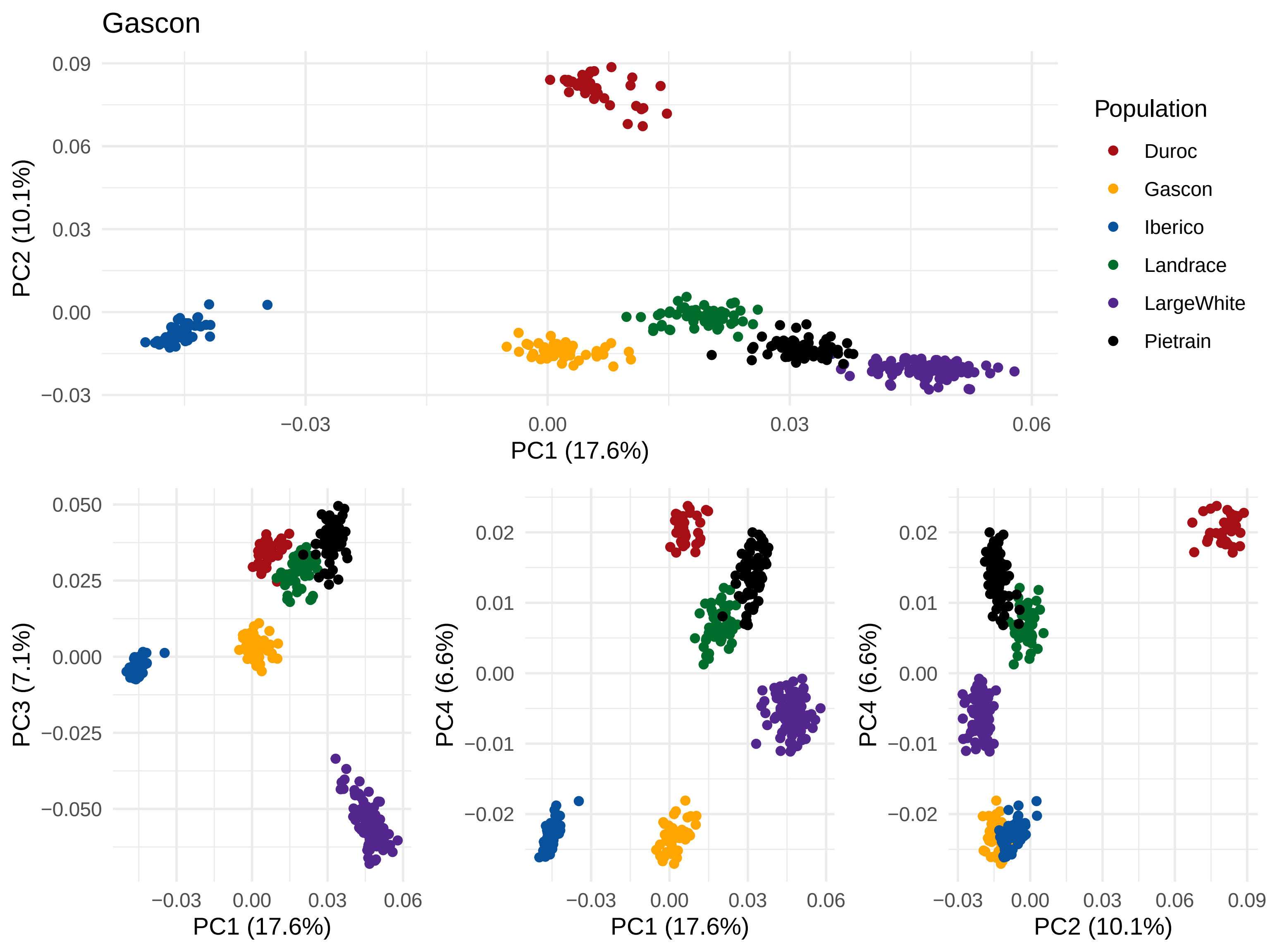

Gascon
Population
Duroc
Gascon
Iberico
Landrace
LargeWhite
Pietrain
PC1 (17.6%)
PC2 (10.1%)
PC3 (7.1%)
PC4 (6.6%)

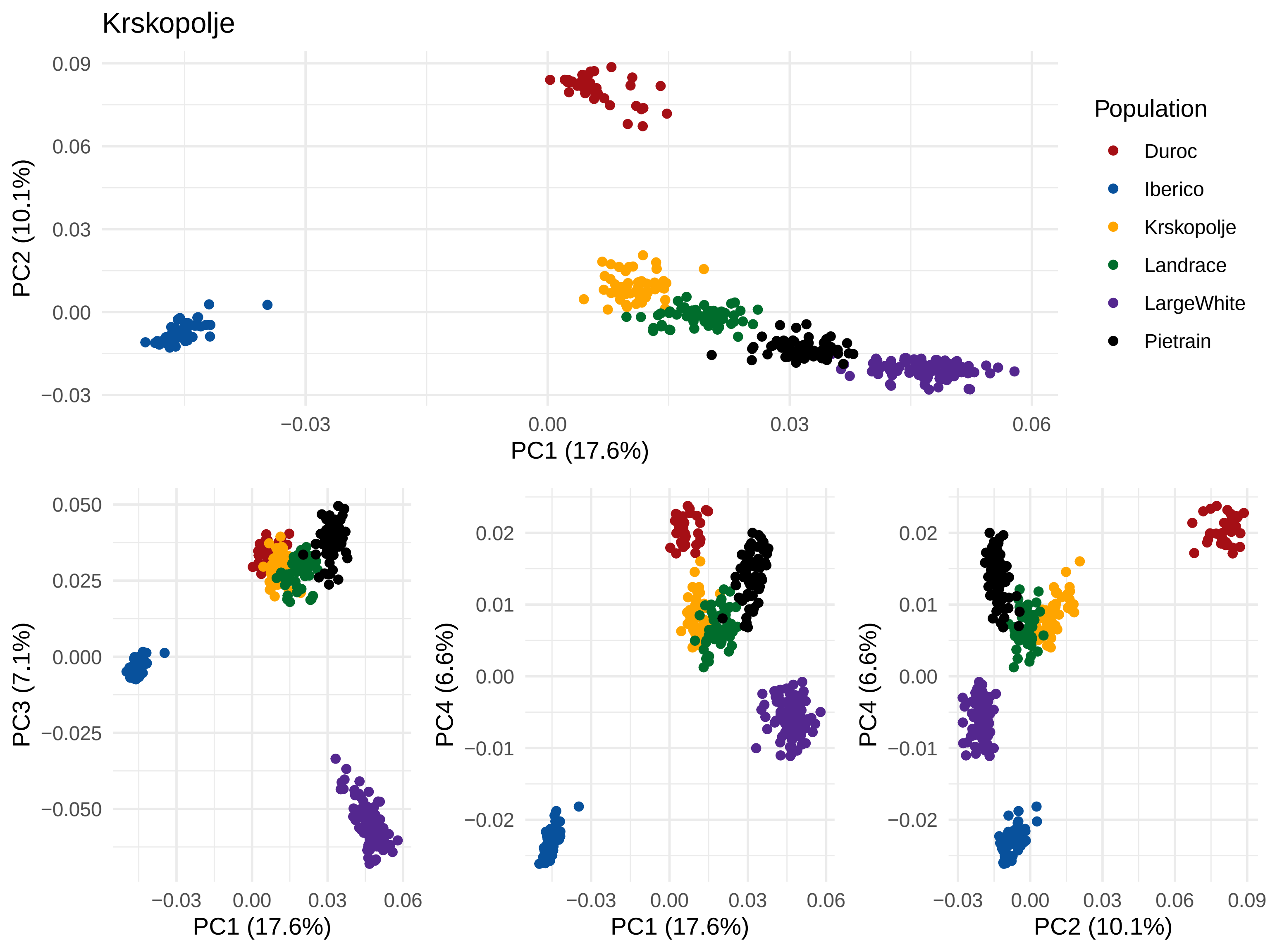
Krskopolje
PC1 (17.6%)
PC2 (10.1%)
0.09
0.06
0.03
0.00
−0.03
−0.03
0.00
0.03
0.06
Population
Duroc
Iberico
Krskopolje
Landrace
LargeWhite
Pietrain
PC3 (7.1%)
0.050
0.025
0.000
−0.025
−0.050
PC1 (17.6%)
PC4 (6.6%)
0.02
0.01
0.00
−0.01
−0.02
PC1 (17.6%)
PC4 (6.6%)
PC2 (10.1%)
0.09

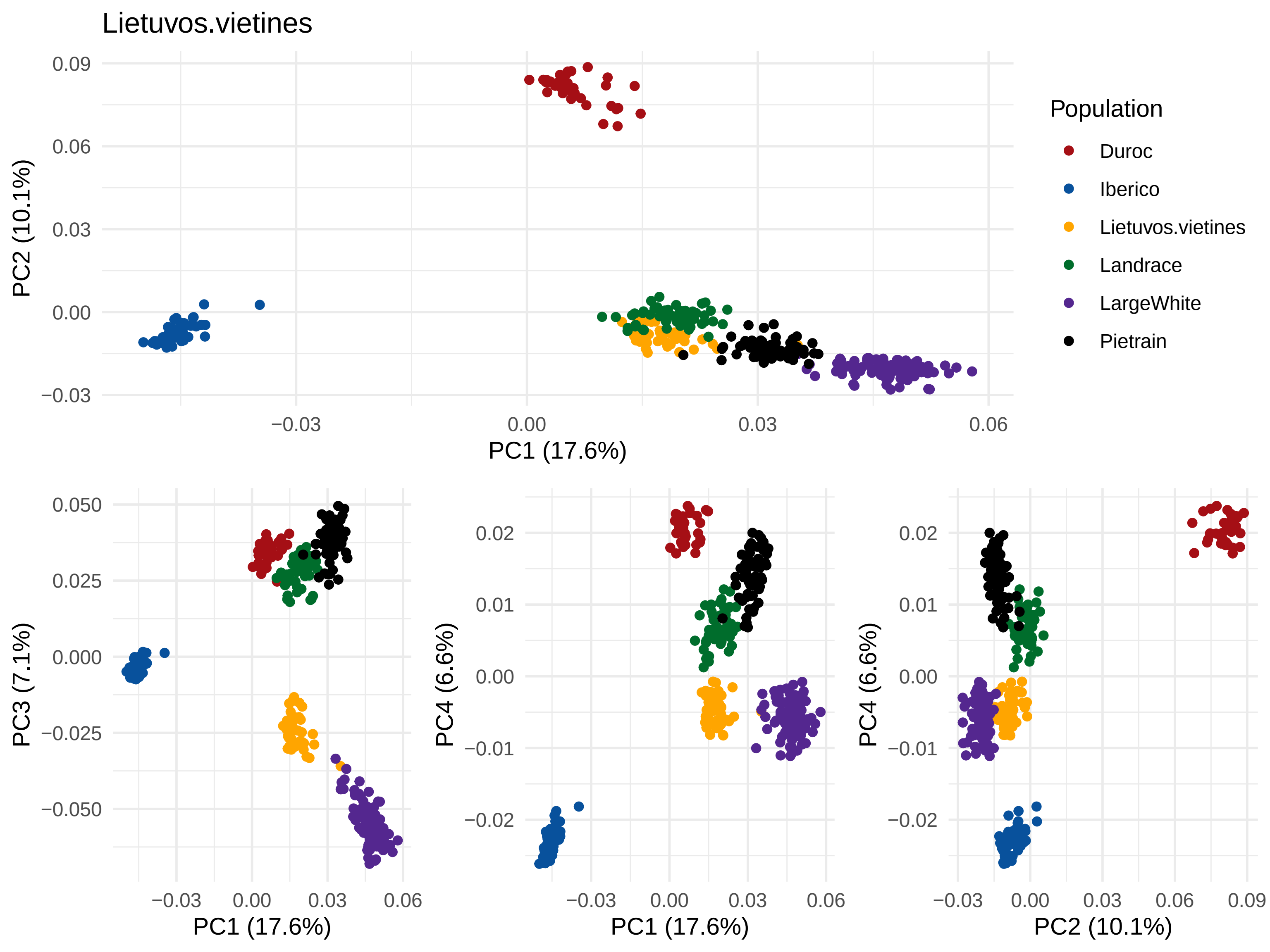
Lietuvos.vietines
PC2 (10.1%)
PC1 (17.6%)
Population
Duroc
Iberico
Lietuvos.vietines
Landrace
LargeWhite
Pietrain
PC3 (7.1%)
PC1 (17.6%)
PC4 (6.6%)
PC1 (17.6%)
PC4 (6.6%)
PC2 (10.1%)

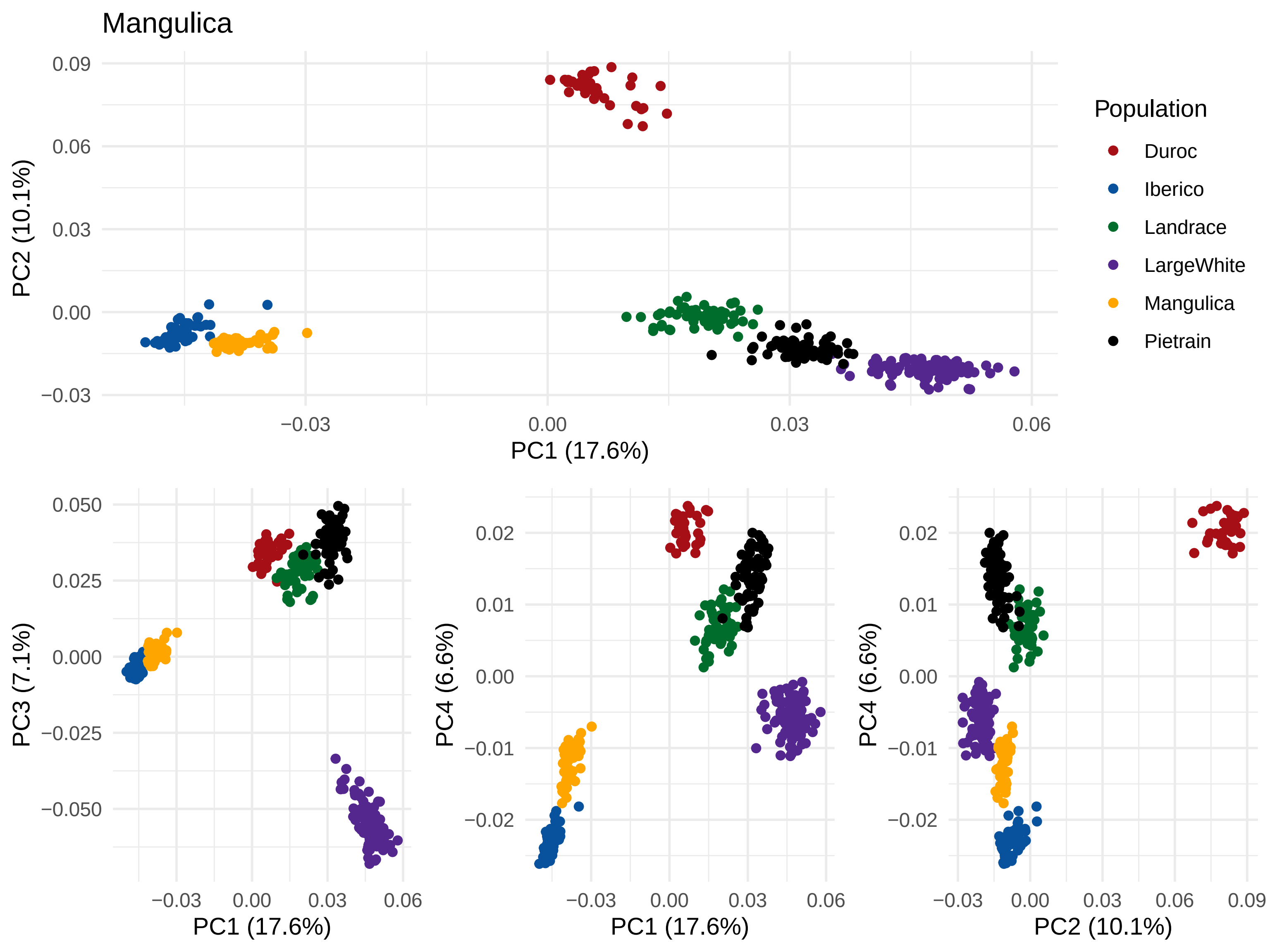
Mangulica
PC2 (10.1%)
PC1 (17.6%)
Population
Duroc
Iberico
Landrace
LargeWhite
Mangulica
Pietrain
PC3 (7.1%)
PC4 (6.6%)
PC2 (10.1%)

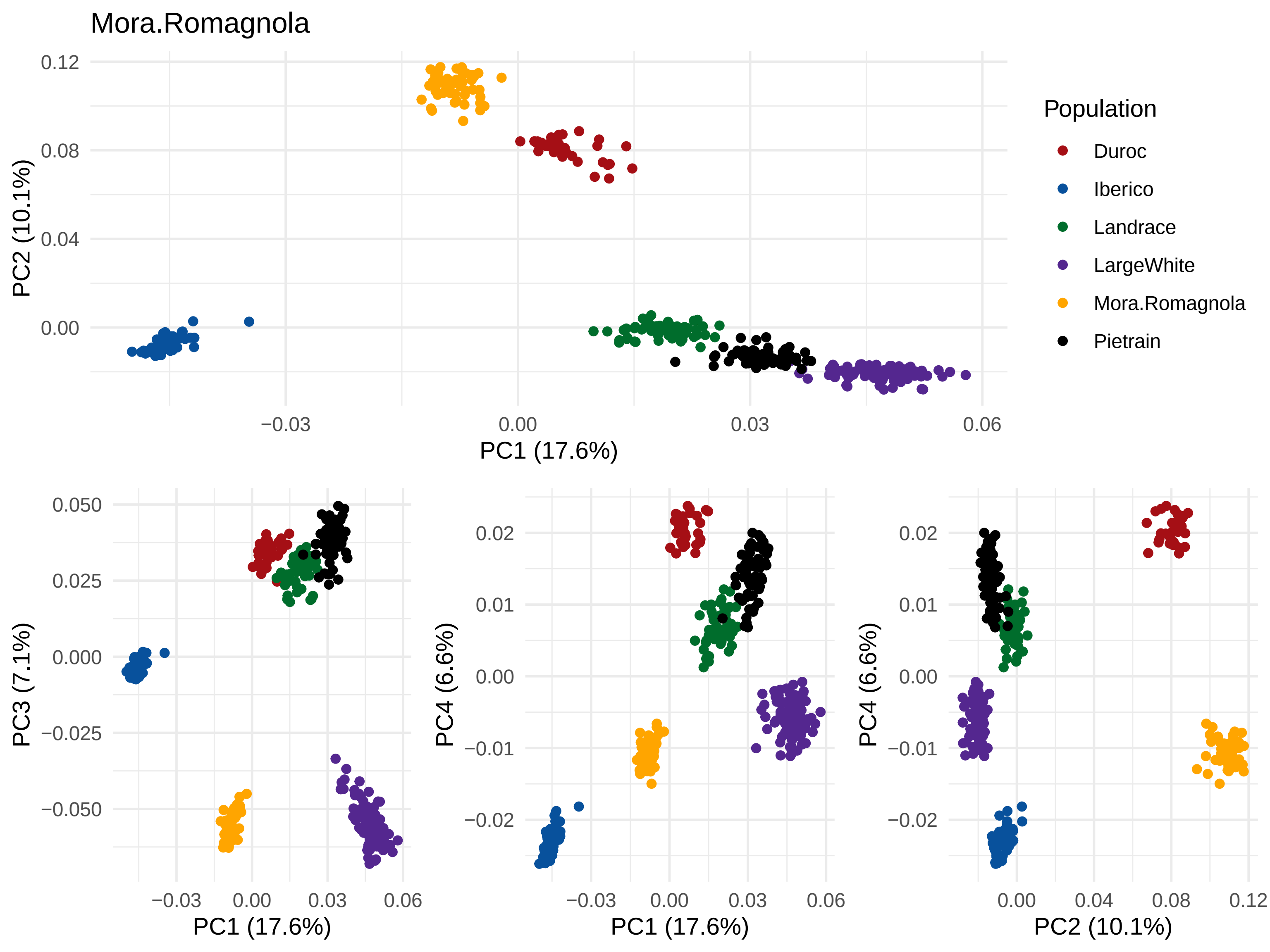
Mora.Romagnola
PC1 (17.6%)
PC2 (10.1%)
PC3 (7.1%)
PC4 (6.6%)
Population
Duroc
Iberico
Landrace
LargeWhite
Mora.Romagnola
Pietrain

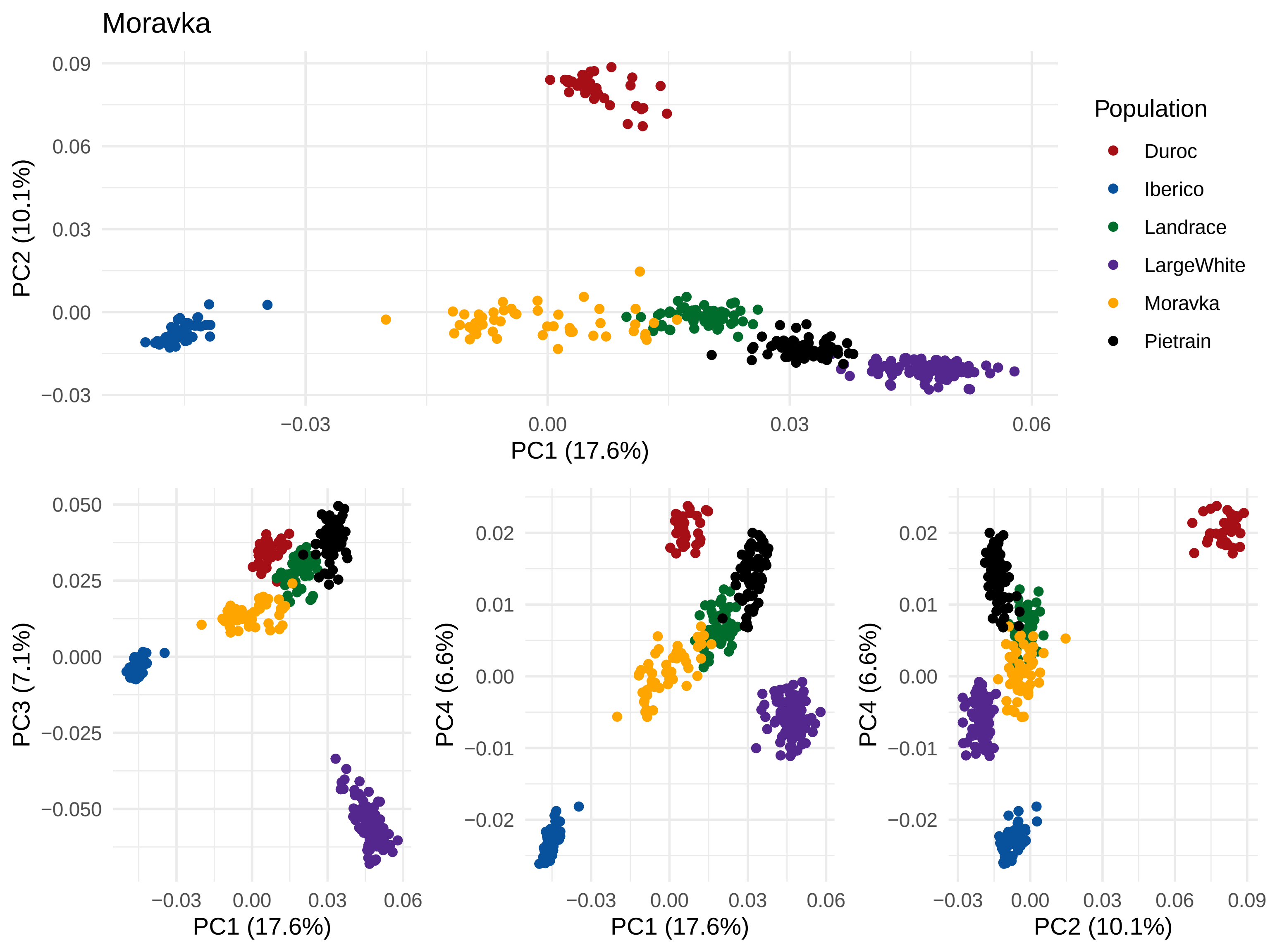

Moravka
PC1 (17.6%)
PC2 (10.1%)
PC3 (7.1%)
PC4 (6.6%)
Population
Duroc
Iberico
Landrace
LargeWhite
Moravka
Pietrain

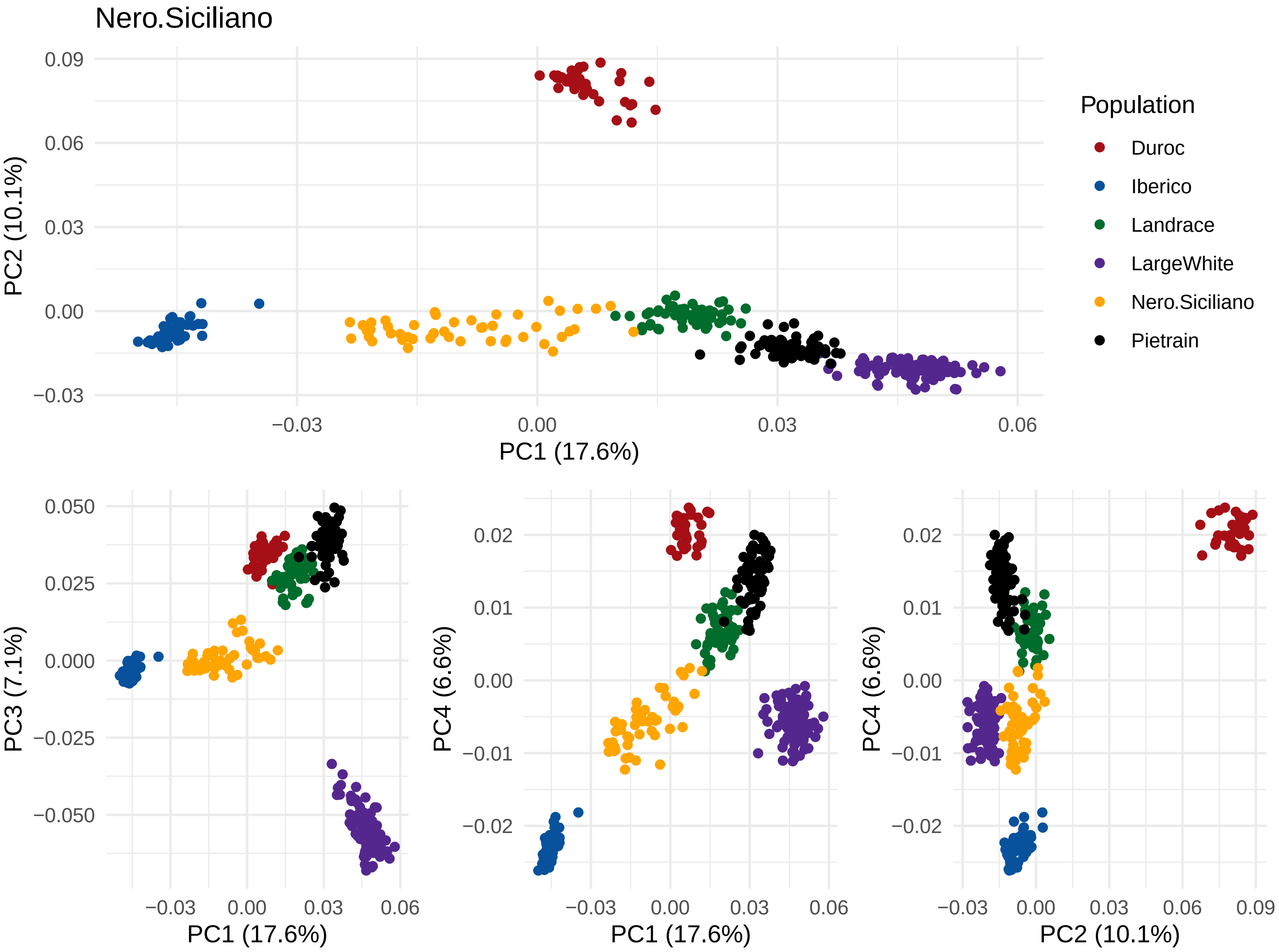

Nero.Siciliano
PC2 (10.1%)
PC1 (17.6%)
0.09
0.06
0.03
0.00
−0.03
−0.03
0.00
0.03
0.06
Population
Duroc
Iberico
Landrace
LargeWhite
Nero.Siciliano
Pietrain
PC3 (7.1%)
0.050
0.025
0.000
−0.025
−0.050
PC4 (6.6%)
0.02
0.01
0.00
−0.01
−0.02
PC2 (10.1%)
0.09

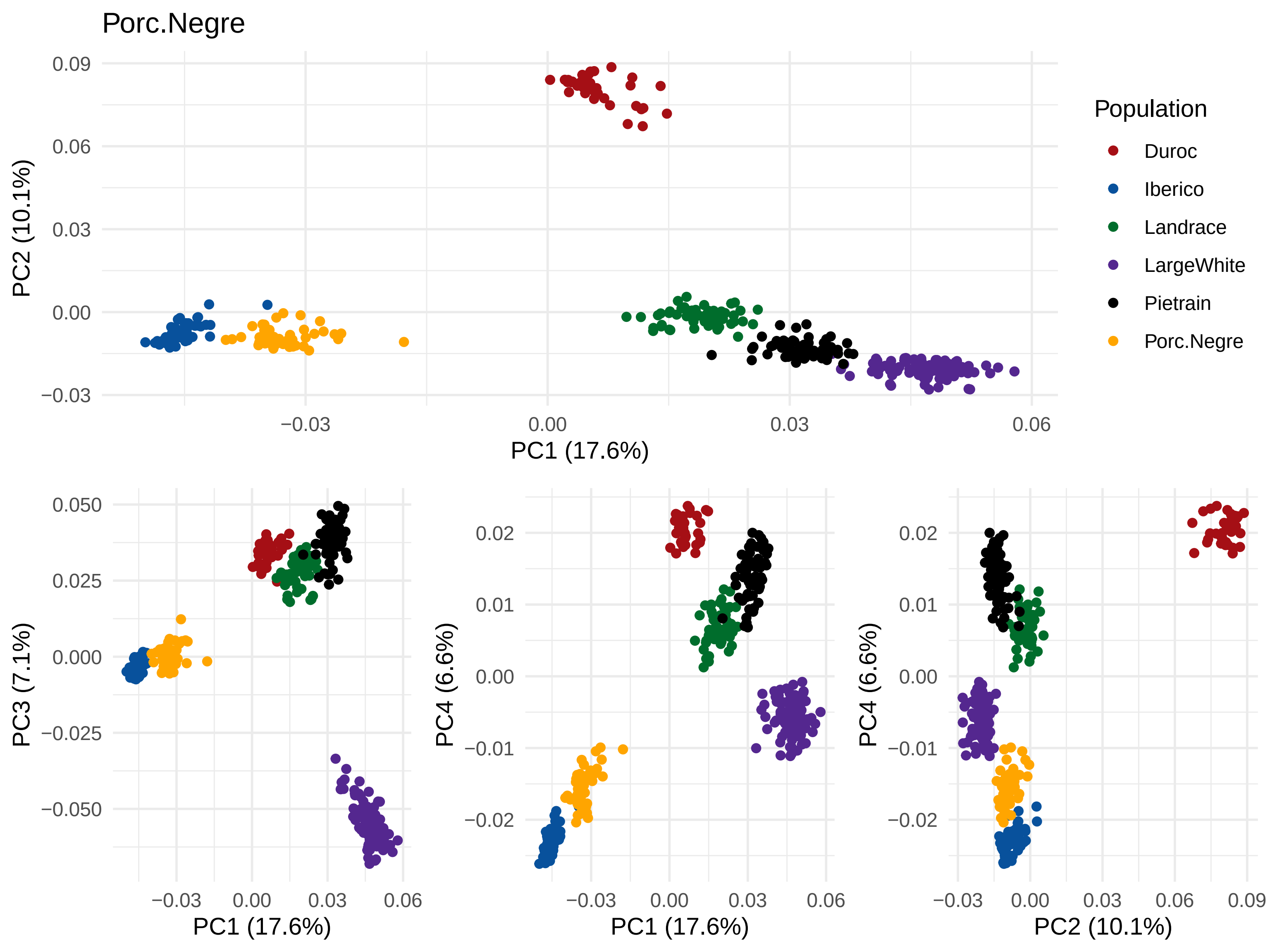
Porc.Negre
PC1 (17.6%)
PC2 (10.1%)
PC3 (7.1%)
PC4 (6.6%)
Population
Duroc
Iberico
Landrace
LargeWhite
Pietrain
Porc.Negre

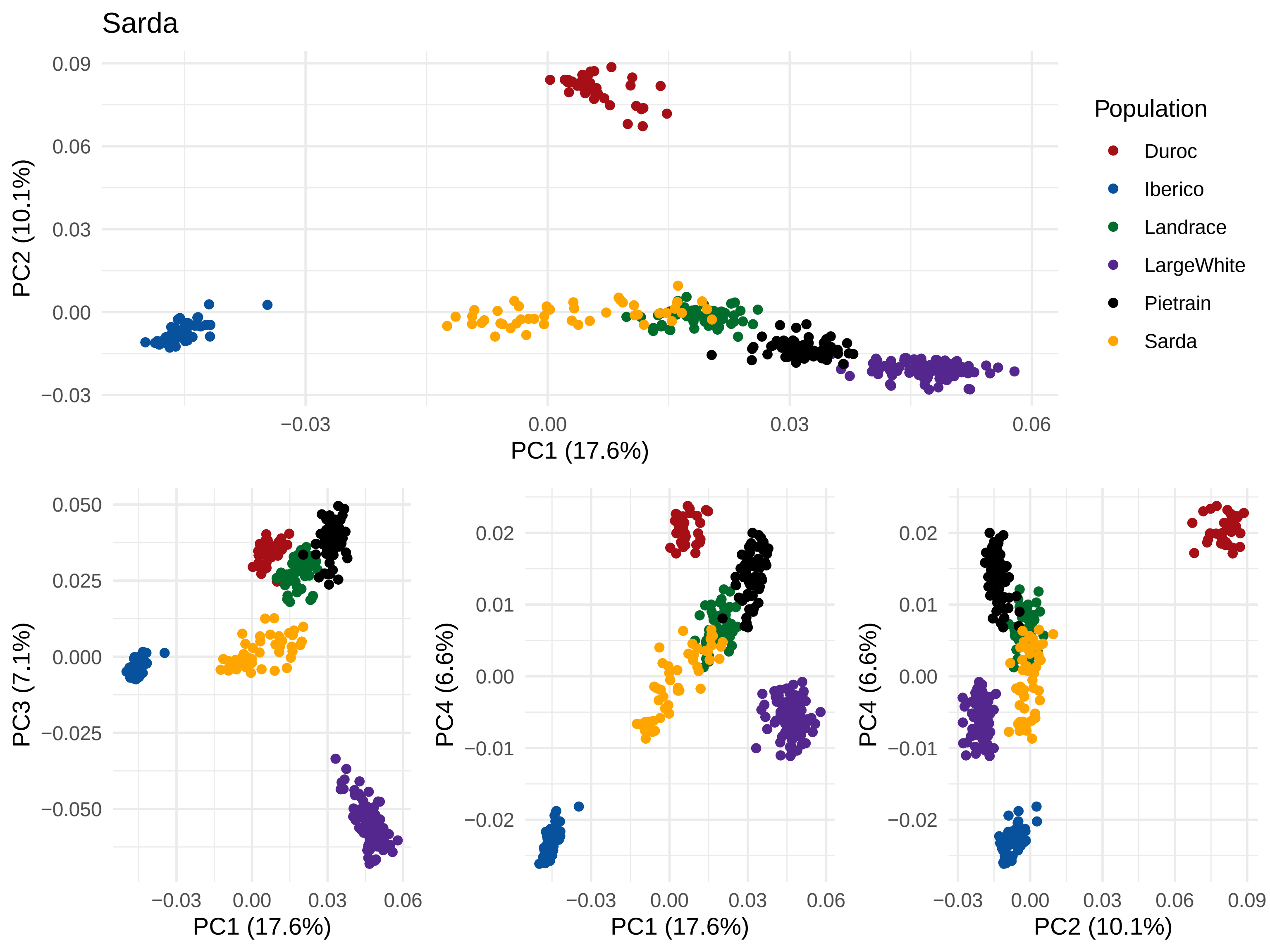

Sarda
PC2 (10.1%)
PC1 (17.6%)
0.09
0.06
0.03
0.00
−0.03
−0.03
0.00
0.03
0.06
Population
Duroc
Iberico
Landrace
LargeWhite
Pietrain
Sarda
PC3 (7.1%)
0.050
0.025
0.000
−0.025
−0.050
PC4 (6.6%)
0.02
0.01
0.00
−0.01
−0.02
0.09

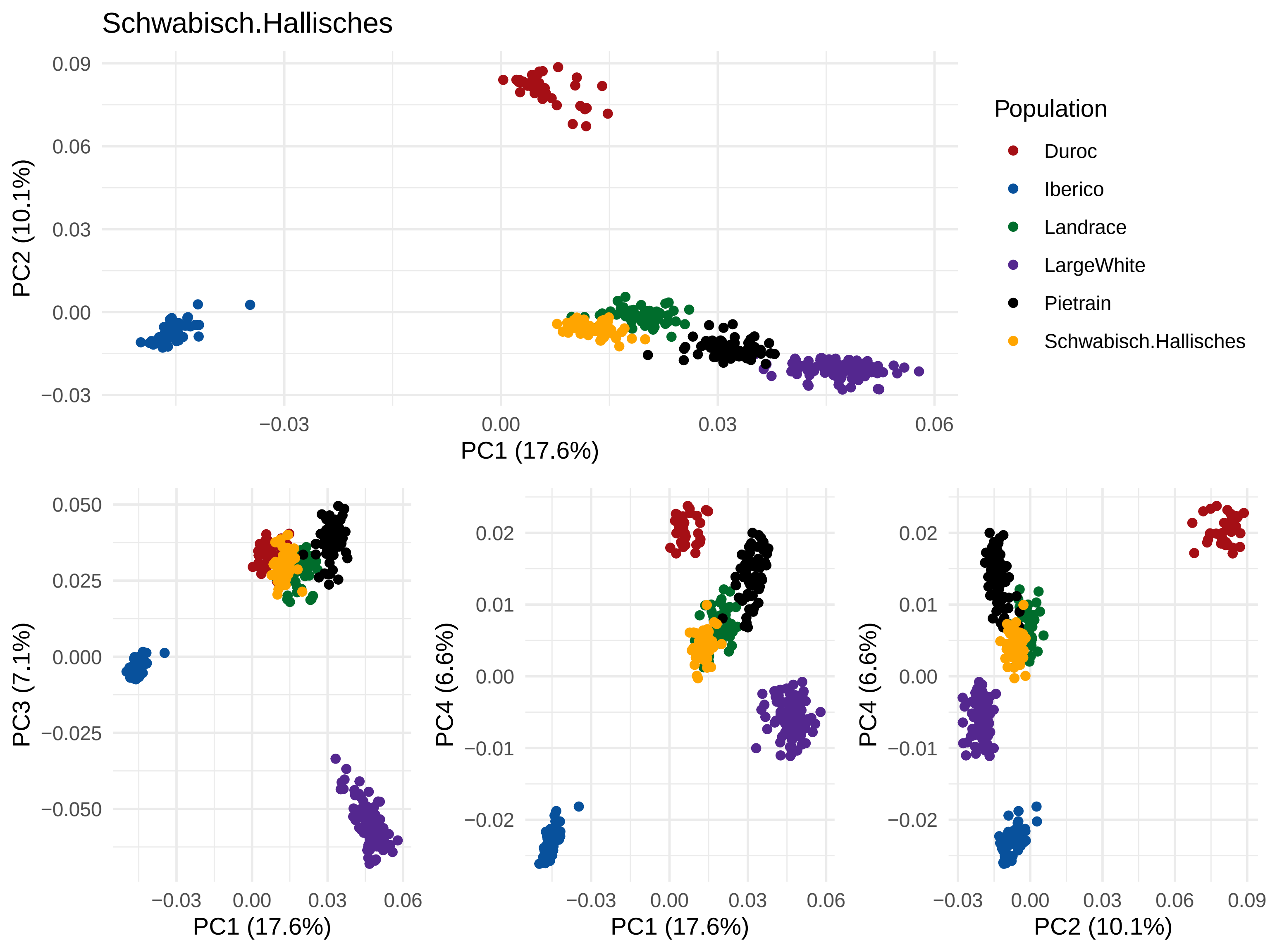
Schwabisch.Hallisches
PC1 (17.6%)
PC2 (10.1%)
PC3 (7.1%)
PC4 (6.6%)
Population
Duroc
Iberico
Landrace
LargeWhite
Pietrain
Schwabisch.Hallisches

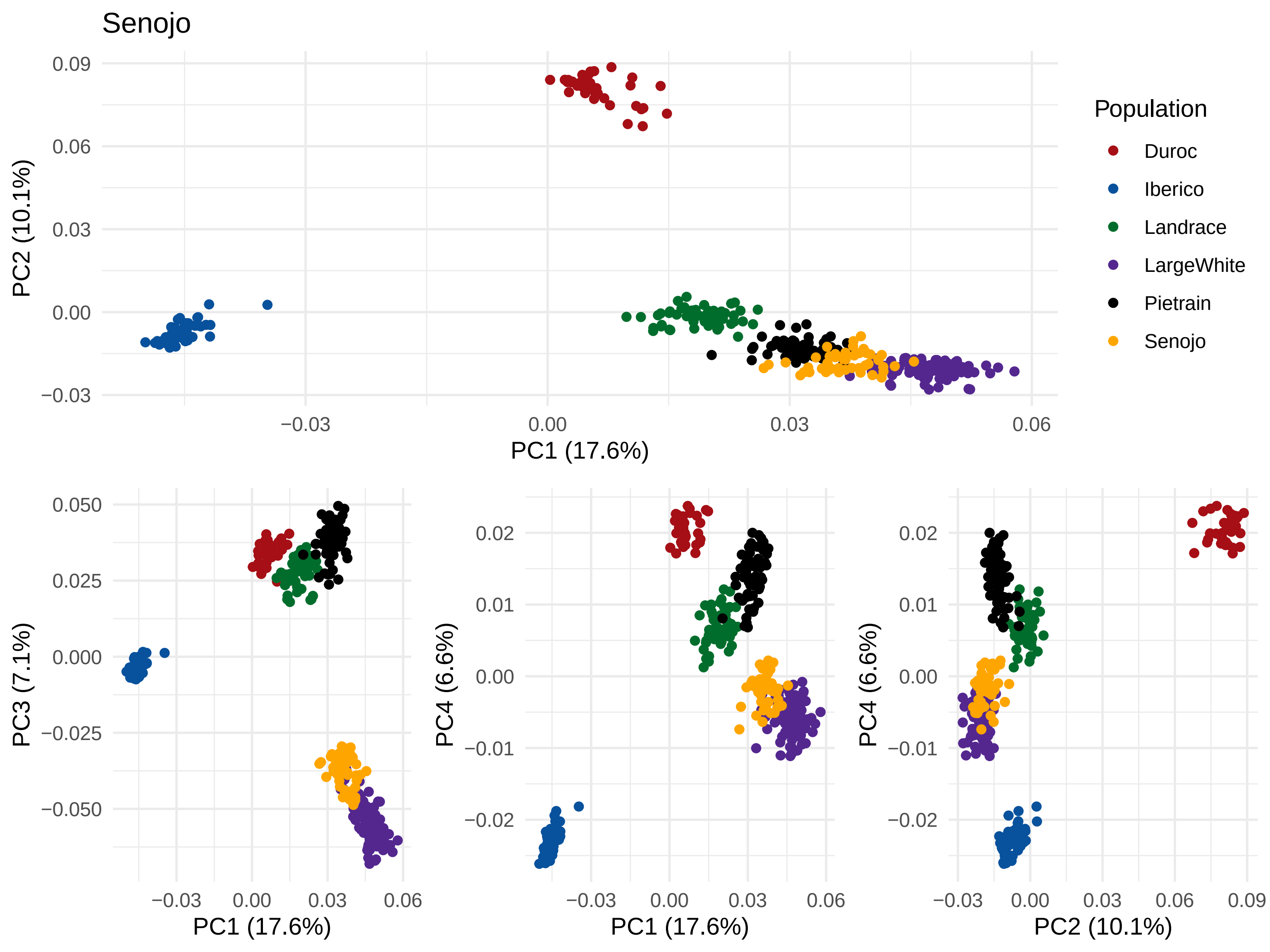

Senojo
PC1 (17.6%)
PC2 (10.1%)
PC3 (7.1%)
PC4 (6.6%)
Population
Duroc
Iberico
Landrace
LargeWhite
Pietrain
Senojo

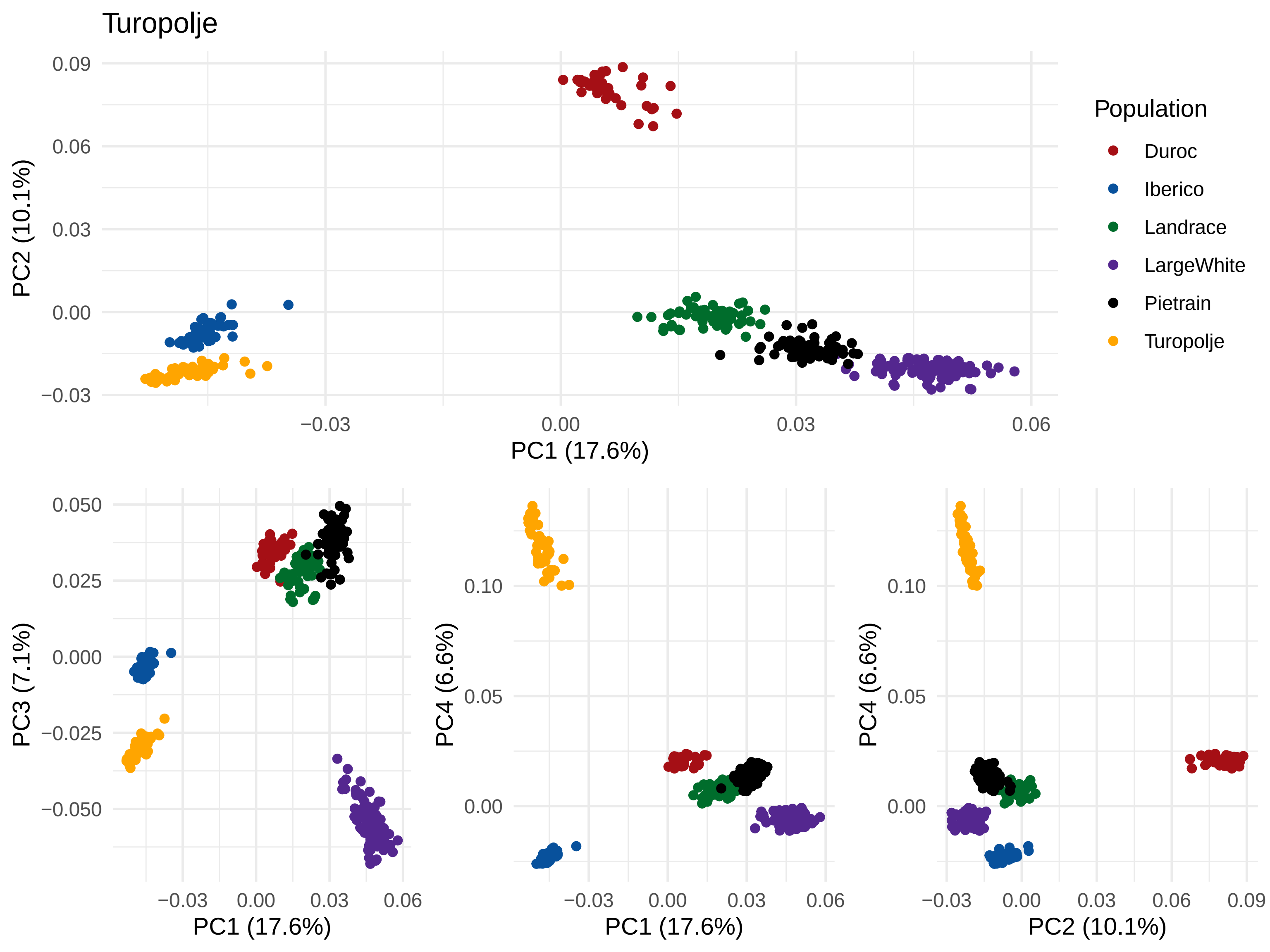
Turopolje
Population
Duroc
Iberico
Landrace
LargeWhite
Pietrain
Turopolje
PC1 (17.6%)
PC2 (10.1%)
PC3 (7.1%)
PC4 (6.6%)
0.09
0.06
0.03
0.00
−0.03
0.050
0.025
0.000
−0.025
−0.050
0.10
0.05
0.00
0.06
0.09
